# Structural Basis for Human Milk Oligosaccharides Recognition by Evolutionarily Dominant Norovirus Genogroup II Genotypes

**DOI:** 10.64898/2026.11.04.764187

**Authors:** Luca Pisapia, Henry N. Flatau, Siyu Lin, Rebecca Calamandrei, Linda Cerofolini, Juan C. Muñoz-García, Anne Imberty, Marco Fragai, Jesus Angulo, Annabelle Varrot

## Abstract

Human noroviruses are a major cause of acute gastroenteritis. Human milk oligosaccharides (HMOs) contain glycan motifs similar to histo-blood group antigens and may act as soluble decoys that reduce viral attachment. However, the molecular basis of HMO recognition by the P domain of the major capsid protein VP1 of different norovirus genotypes is not fully understood. Here, we combined ligand-based and protein-based NMR experiments with X-ray crystallography to study HMO binding to the P domain of the latent GII.2, dominant GII.4 and emerging GII.17 noroviruses genotypes. Screening of 12 HMOs by STD NMR showed binding of all fucosylated HMOs to GII.4 and GII.17 and none for non-fucosylated HMOs. In contrast, GII.2 showed little or no detectable binding under the same conditions. Seven crystal structures of GII.4 or GII.17 with complexed HMOs supported fucose anchoring via five conserved residues. The remaining sugar moieties formed genotype-specific secondary interactions with an extended, shallow and solvent exposed binding surface whose shape is dictated by divergent surface loops. GII.4 recognition is mainly driven by hydrogen bonds whereas GII.17 relied more strongly on water-mediated interactions. Chemical shift perturbation from NMR titrations (CSP NMR) of GII.4 corroborated the fucose interactions and gave dissociation constants of 20-33 mM, consistent with weak and rapidly exchanging interactions.

**IMPORTANCE:** This work highlights the potential of HMOs as decoy receptors against norovirus infections. Further, it provides structural information on the broad binding profile of P domain to support the future development of HMOs derivatives or glycan-based molecules to inhibit noroviruses attachment.

## INTRODUCTION

Human noroviruses (HuNoVs) are the leading infectious cause of acute non-bacterial gastroenteritis worldwide. They are responsible for an estimated 685 million episodes and approximately 212,000 deaths annually across all age groups, second only to rotavirus among enteric pathogens, exposing infants and elderly at great risks (1, 2). However, no approved antivirals nor licensed vaccines are currently available, highlighting a substantial unmet therapeutic need (2).

Noroviruses are classified into ten genogroups (GI–GX), each further subdivided into genotypes based on the VP1 capsid amino acid sequence divergence and into P-types based on nucleotide diversity in the RNA-dependent RNA polymerase (RdRp) region, yielding at least 49 genotypes and 60 P-types across the genus (3). Genogroups I and II are responsible for the overwhelming majority of human infections. Within genogroup II, genotype 4 (GII.4) has dominated global norovirus epidemiology since the mid-1990s through the periodic emergence of antigenically distinct variants that accumulate mutations in the surface-exposed P2 subdomain of the major capsid protein VP1 and thereby achieve partial escape from pre-existing immunity (4, 5). Starting in 2014, GII.17 surged across Asia and displaced GII.4 as the dominant genotype in several East Asian countries (6, 7), while a recombinant GII.2 lineage accounted for up to 20% of reported outbreaks in 2016–2017 before subsiding (8, 9). The lasting co-circulation of these genetically distinct lineages makes a comparative understanding of host receptor engagement across multiple HuNoVs strains an important goal.

The susceptibility of humans to norovirus infections was first linked to histo-blood group antigen (HBGA) status in early studies demonstrating that virus-like particles bind these carbohydrate antigens in a strain-dependent manner (10, 11). HBGAs are structurally diverse fucosylated oligosaccharides expressed on the surface of red blood cells and on the vascular endothelium of all individuals. Additionally, they are present in secretions such as saliva or intestinal mucus in approximately 80% of the population, designated as secretor-positive individuals. The ABH and Lewis systems define the principal antigen families that differ in the nature and linkage of their terminal fucose and galactose modifications (12). Host susceptibility is strongly influenced by secretor status, as individuals lacking a functional FUT2 α(1,2)-fucosyltransferase are broadly resistant to infection by the most prevalent strains (13), highlighting the importance of the fucose in the infection.

The norovirus capsid is assembled from 180 copies of VP1 with T=3 icosahedral symmetry. VP1 folds into a shell (S) domain and a protruding (P) domain from which the distal P2 subdomain carries the HBGA-binding site and the principal neutralizing epitopes (**Fig. 1**) (14, 15). The P domain assembles as a homodimer, which is the form commonly used to study binding interactions. Crystal structures of GII.4, GII.10 and GII.17 P domain dimers in complex with HBGA oligosaccharides have defined a conserved binding site at the P2 subdomain dimer interface (15, 16). In those genotypes, α-fucose is recognized through a conserved scaffold comprising an invariant threonine, arginine and aspartate from one protomer and an invariant glycine and tyrosine from the other protomer of the dimer. Dominant interactions involve the side chains of arginine and aspartate, supported by main-chain contacts from threonine (where mutations may be tolerated due to the invariant nature of main-chain interactions) and glycine, while tyrosine is involved in a CH-π contact. Secondary interactions with the remaining sugar units vary between genotypes and strains (15–18). The genotype GII.2 was classified originally as an HBGA non-binder nevertheless binding was then observed on A, B and AB type saliva and not on synthetic oligosaccharides (19, 20). Although no crystal structure in complex with a carbohydrate ligand has been reported to date, the apo structures of two variants reveal an equivalent scaffold for fucose recognition (19, 20).

**Figure 1.**
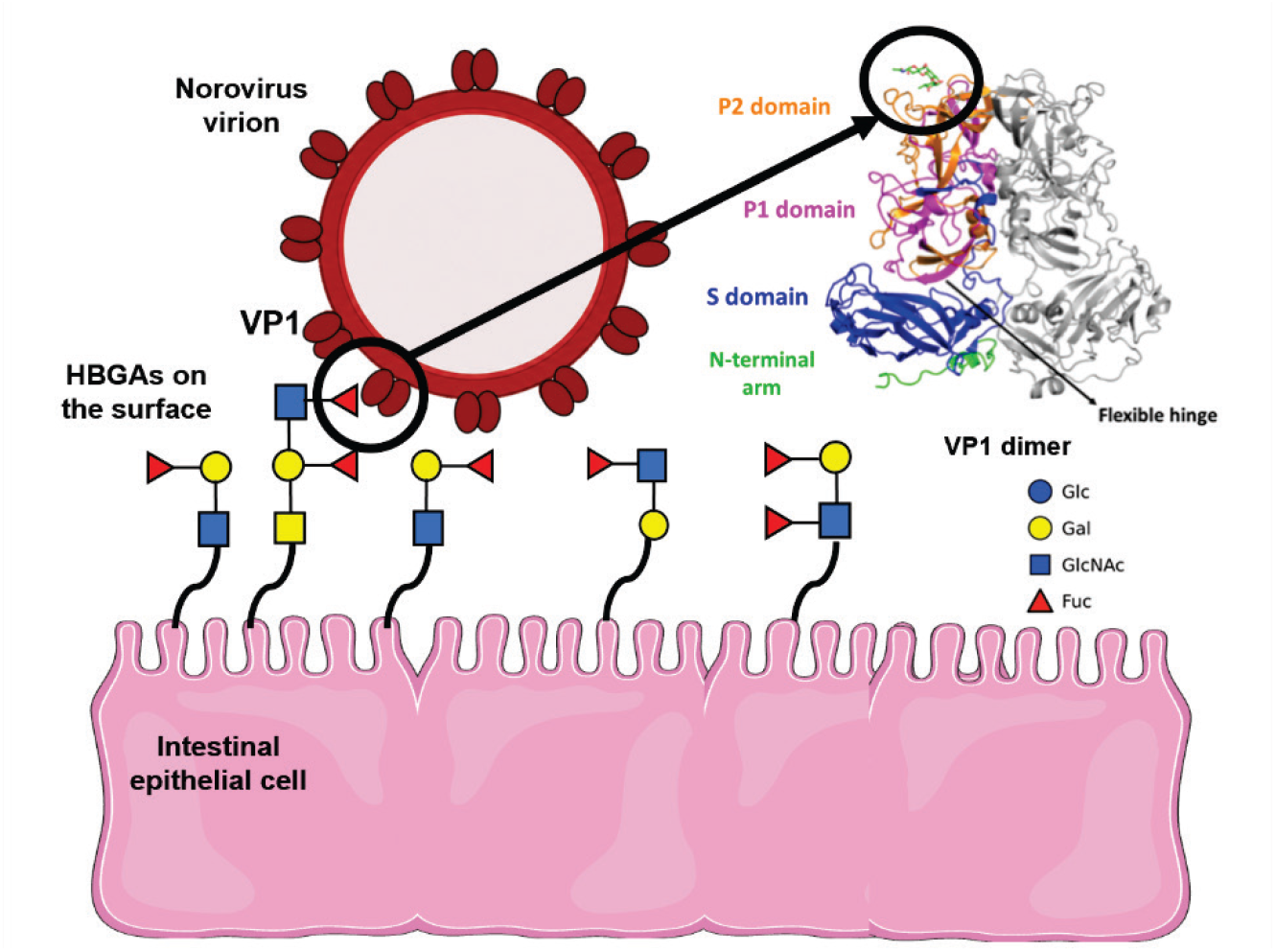
P domain organization of norovirus VP1 and engagement with host histo-blood group antigens (HBGAs). The norovirus particle is composed of 180 copies of VP1 assembled as dimers and organized into N-terminal arm (green), a shell domain (S, blue), a flexible hinge, P1 subdomain (magenta) and distal P2 subdomain (orange), which carries the HBGA-binding site. HBGAs are depicted using SNFG notation (21). VP1 dimer structure from the GI.1 Norwalk virus capsid (PDB 1IHM) (14).

The same fucosylated epitopes that mediate cell-surface HBGA recognition are structurally mimicked in human milk oligosaccharides (HMOs). HMOs are the third most abundant solid component of breast milk, present at concentrations of 5–20 g/L (22). Many HMOs carry α1-2 or α1-3 linked fucose on a lactose core or to a lacto-N-tetraose backbone, generating structural epitopes that mimic secretor and Lewis blood group antigens. Therefore, it was proposed that HMOs act as soluble receptor decoys competing with cell-surface HBGAs for norovirus binding (22–24). X-ray crystallographic studies confirmed that 2′-fucosyllactose (2′FL) and 3-fucosyllactose (3FL) occupy the HBGA-binding site on GII.4, GII.10 and GII.17 P domain dimers, reproducing key contacts made by H-type antigens and blocking VLP attachment to HBGAs in a dose-dependent manner (25, 26). These analyses were, however, restricted to simple trisaccharide species and to individual genotypes. The binding mode of the more complex branched fucosylated HMOs that predominate in milk from secretor-positive women, such as lacto-N-fucopentaoses I, II and III (LNFP-I, -II, -III), lacto-N-difucohexaose I (LNDFH-I), difucosyllactose (DFL) and fucosyl-sialyllactose (FSL) has not been characterized for any human genotype (**Fig. 2**).

**Figure 2.**
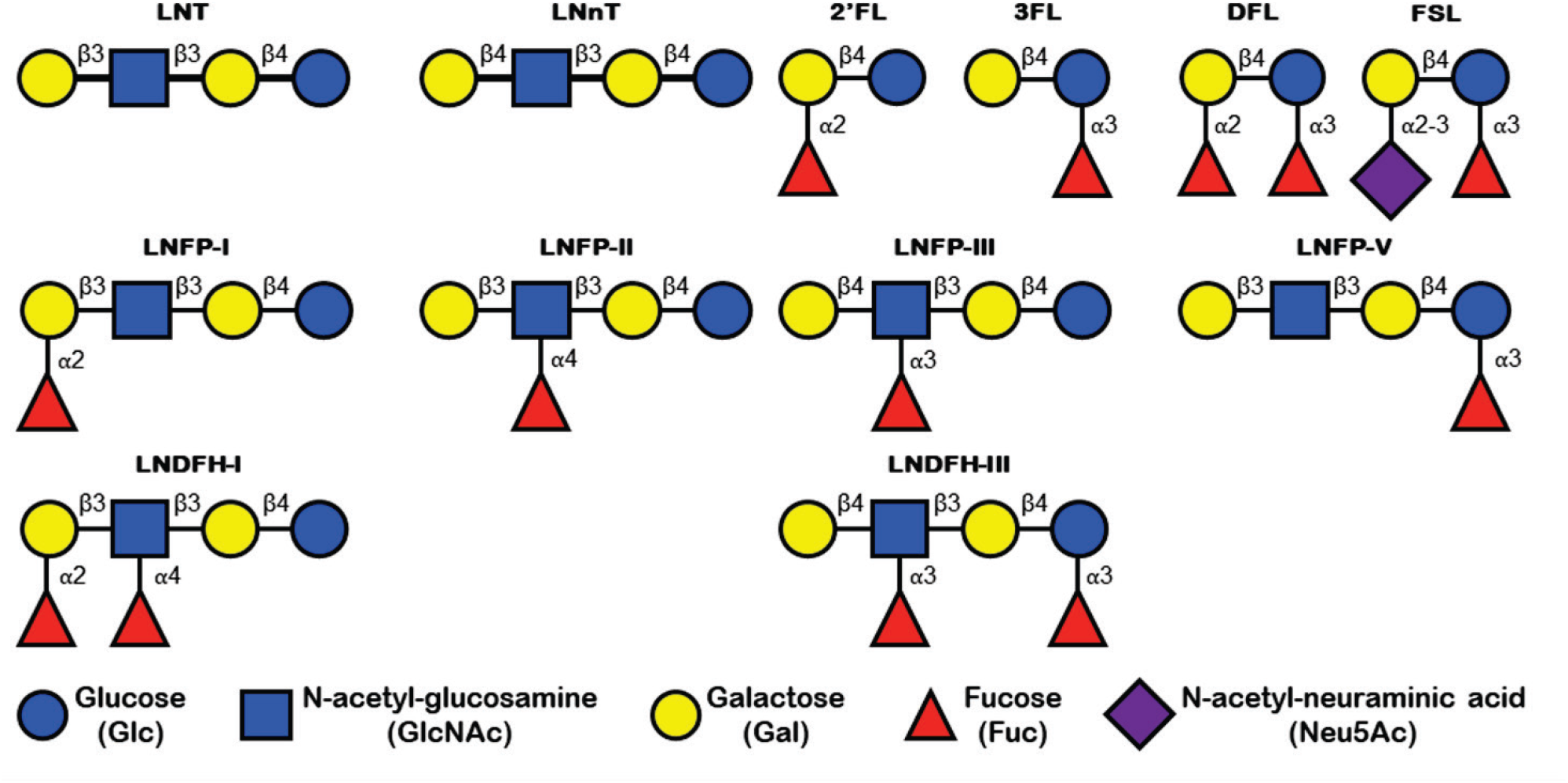
Description of the HMOs used for screening using the SNFG convention. X-ray crystallography on recombinant P domain co-crystals has been an invaluable tool to characterize norovirus–carbohydrate interactions at the atomic level, defining the hydrogen-bonding networks, water-mediated contacts and hydrophobic interactions governing recognition (15–18). Saturation transfer difference (STD) NMR spectroscopy provides a quick solution for rapid screening for interactions, while also providing a solution-state view of the same interactions by mapping which ligand protons are in closest contact with the protein surface in the transiently bound complex (binding epitope maps) (27, 28). NMR titrations of isotopically enriched P domain dimers monitor chemical shift perturbations (CSP) and enable analysis of binding-induced changes at the protein side. Applied in combination, the three approaches provide structural, epitope and thermodynamic information that are mutually reinforcing and experimentally non-redundant.

Here, we report a combined study of the interactions involving the recombinant P domain of three circulating strains: latent GII.2/US/2022 (GII.2), the dominant GII.4/Sydney/2016 (GII.4) and emerging GII.17/Rotterdam/2024 (GII.17) with a panel of 12 naturally occurring HMOs (**Fig. 2**). The screening by STD NMR revealed interactions with every fucosylated HMO for all three genotypes whilst the crystallization screening led to seven crystal structures of GII.4 and GII.17 P domain dimers in complex with fucosylated HMOs of varying length (from tetra-to hexasaccharide) at high resolution, but none for GII.2. Alongside the conserved fucose binding site, uncovered genotype-specific secondary recognition surfaces were revealed. STD NMR binding epitope maps and CSP NMR affinity measurements validate the crystallographic contacts in solution and confirm low millimolar affinity. Together, these findings broaden the current perspective of HMOs as decoy receptors against norovirus infection by demonstrating that, beyond short HMOs, longer HMOs also engage the viral P domain.

## RESULTS

### STD NMR screening of Human Milk Oligosaccharides against NoV

An HMO library containing a total of 12 natural HMOs, provided by Glycom A/S, was first screened against the three circulating NoV genotypes: GII.2 USA, GII.4 Sydney and GII.17 Rotterdam by STD NMR spectroscopy (**Fig. 2**). This method allows to be near physiological conditions in solution and to detect binding while it also makes ligand binding epitope mapping possible, thereby reporting for each ligand the main “hot spots” for the interaction with the protein. Successful binding events were easily monitored by the detection of the STD NMR signal of the methyl protons of fucose. For LNT and LNnT, which do not contain fucose, a binding event was monitored by the observation of the anomeric protons STD NMR signals.

For the GII.4 and GII.17 genotypes, binding was detected for 10 out of the 12 HMOs as well as for the monosaccharide L-fucose (**Fig. 3**). Every HMO containing at least one fucose residue showed STD signals, confirming binding whilst the non-fucosylated ligands, LNT and LNnT, displayed no signals beyond the N-acetyl of GlcNAc. This peak is attributed to unspecific binding, due to the absence of interactions from other protons. However, while most ligands showed binding, their very low STD intensities (STD factors from 0.5 to 2.6 %), even at long saturation times (6 s), suggest extremely weak interactions. In general, while clear interactions were observed for the fucose methyl groups (≍ 0.9 ppm) with GII.4 and GII.17, additional interactions were also detected for the anomeric region of the oligosaccharides (4.4 to 5.2 ppm), as well as for the protons around 3.5 ppm (H3s, H4s and H6s and, in some HMOs, also H2s and H5s). However, making clear statements about which of those protons are interacting was challenging, due to overlapping, making it impossible to quantify the STD signal with high confidence (**Fig. S1**).

**Figure 3.**
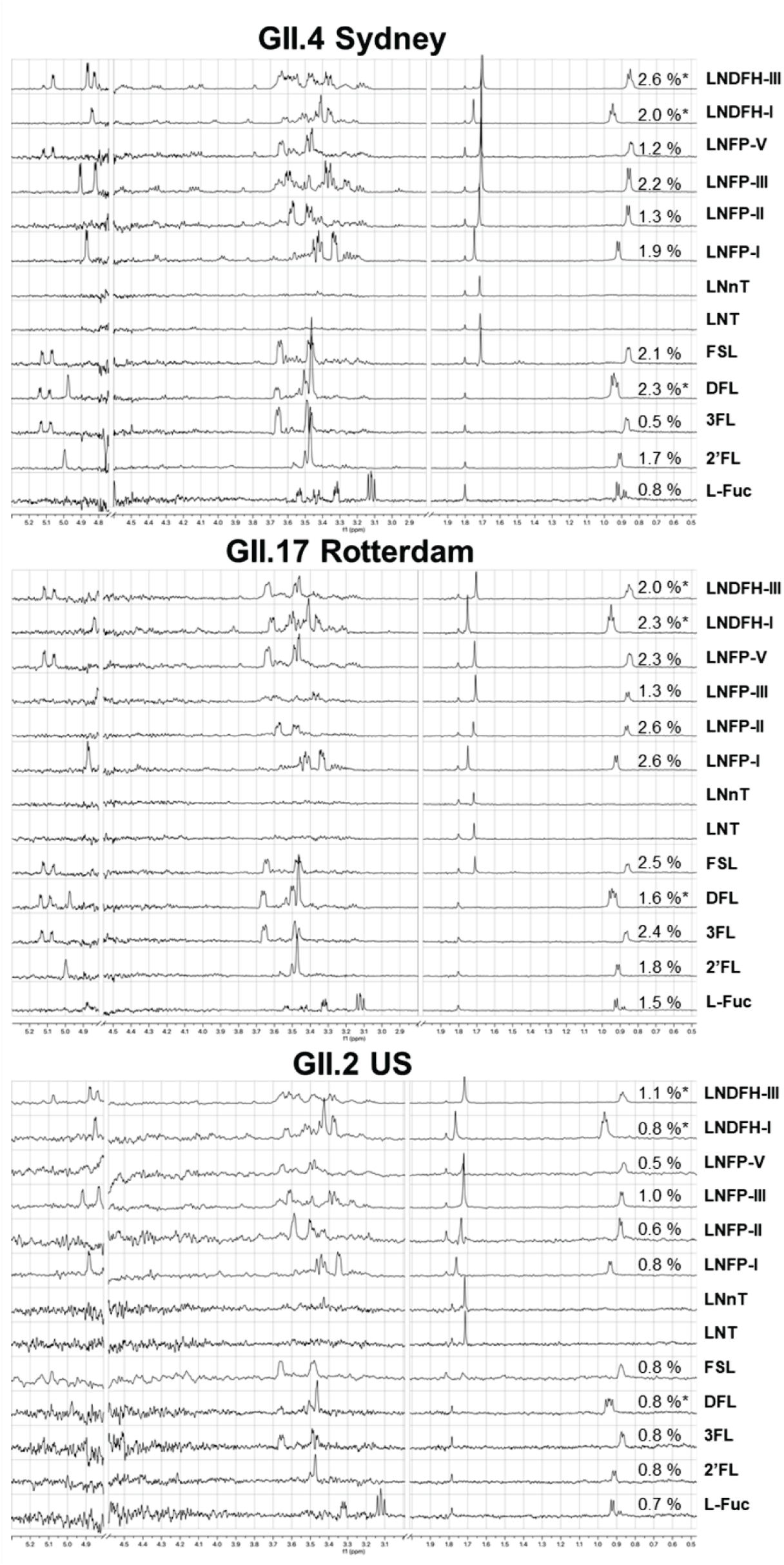
STD NMR spectra (6 s saturation time, 256 scans) of the HMOs screening against the P-dimer of HuNoVs genotypes: GII.2 USA, GII.4 Sydney and GII.17 Rotterdam. The percentages correspond to the STD intensities of the fucose methyl protons (∼ 0.9 ppm), while the asterisk indicates overlapping methyl peaks of two fucoses. Experiments were performed in PBS D_2_O pH 7.4 buffer, at 5 ℃, using 15 µM of protein and a protein-to-ligand ratio of 1:100.

In the screening against GII.17, no anomeric interactions were detectable for LNFP-I, LNFP-II, LNFP-III and LNDFH-I, while LNFP-V and LNDFH-III presented detectable interactions with the anomeric protons. Both LNFP-V and LNDFH-III, bear an *a*(1–3)-L-fucose (*a*3Fuc) branching off the glucose moiety of the lactose core. In contrast, LNDFH-I which also presents an *a*4Fuc, showed no sign of anomeric proton involvement on binding. These results indicate different binding modes for the different HMOs (**Fig. 3**).

Further differences in the STD intensities of HMOs among the different genotypes were observable on the fucose methyl: for the fast kinetics that characterize these interactions, the very low STD intensities of 3FL and L-Fuc are supportive of weak binding interactions to GII.4 (0.5% and 0.8% STD factors, respectively for the methyl protons of Fuc) while larger STD intensities are indicative of stronger binding to GII.17 (2.4 % and 1.5 %, respectively). The stronger preference of GII.17 for the methyl of *a*3Fuc on the reducing end is also shown for LNFP-V, which presents this motif, where the *a*3Fuc methyl showed stronger STD to GII.17 (2.3%) than to GII.4 (1.2%). On the contrary, LNFP-III, which presents an *a*3Fuc on the non-reducing end, showed a higher STD intensity for GII.4 (2.2%) than for GII.17 (1.3%).

In contrast to the GII.4 and GII.17 genotypes, the screening against GII.2 showed, for most protons, STD intensities below 1% at 6 s saturation time, leading to very noisy spectra. This indicates that the interaction with the GII.2 protein is close to the detection limit of the STD NMR technique and of extremely low affinity (**Fig. 3**).

Overall, STD NMR screening clearly demonstrates that the fucose moiety is the key structural requirement for HMOs recognition by HuNoVs GII.4 and GII.17. These results, together with the observation of ligand contacts beyond the fucose moiety for the HMOs, show that sugar moieties of the carbohydrate chain other than fucose also play a role in HMO binding to HuNoVs.

### Binding epitope maps of HMOs to HuNoV GII.4 P-dimers

Due to ^1^H signal overlap, especially in the region around 3.5 ppm, it was not possible to accurately disentangle the STD intensities of each of the individual HMO protons and, hence, generate complete ligand binding epitope maps. Spectral assignment and integration of protons was further complicated due to presence of the *a* and β anomers of each HMO. Despite these challenges, binding epitope maps could be generated for the most abundant HMOs (2’FL, LNFP-III and LNDFH-III) using the reduced dataset-STD NMR (rd-STD NMR) approach **(Fig. 4)**. In structural terms, unlike 2’FL, which resembles closely H-type 2 antigen, LNFP-III resembles at its non-reducing end a Lewis X type HBGA and is also two residues longer than 2’FL. LNDFH-III presents an additional *a*3Fuc on the reducing glucose, making it an interesting ligand to evaluate the impact of an additional fucose residue on binding to the P-dimer as well on potential fucose selectivity.

**Figure 4.**
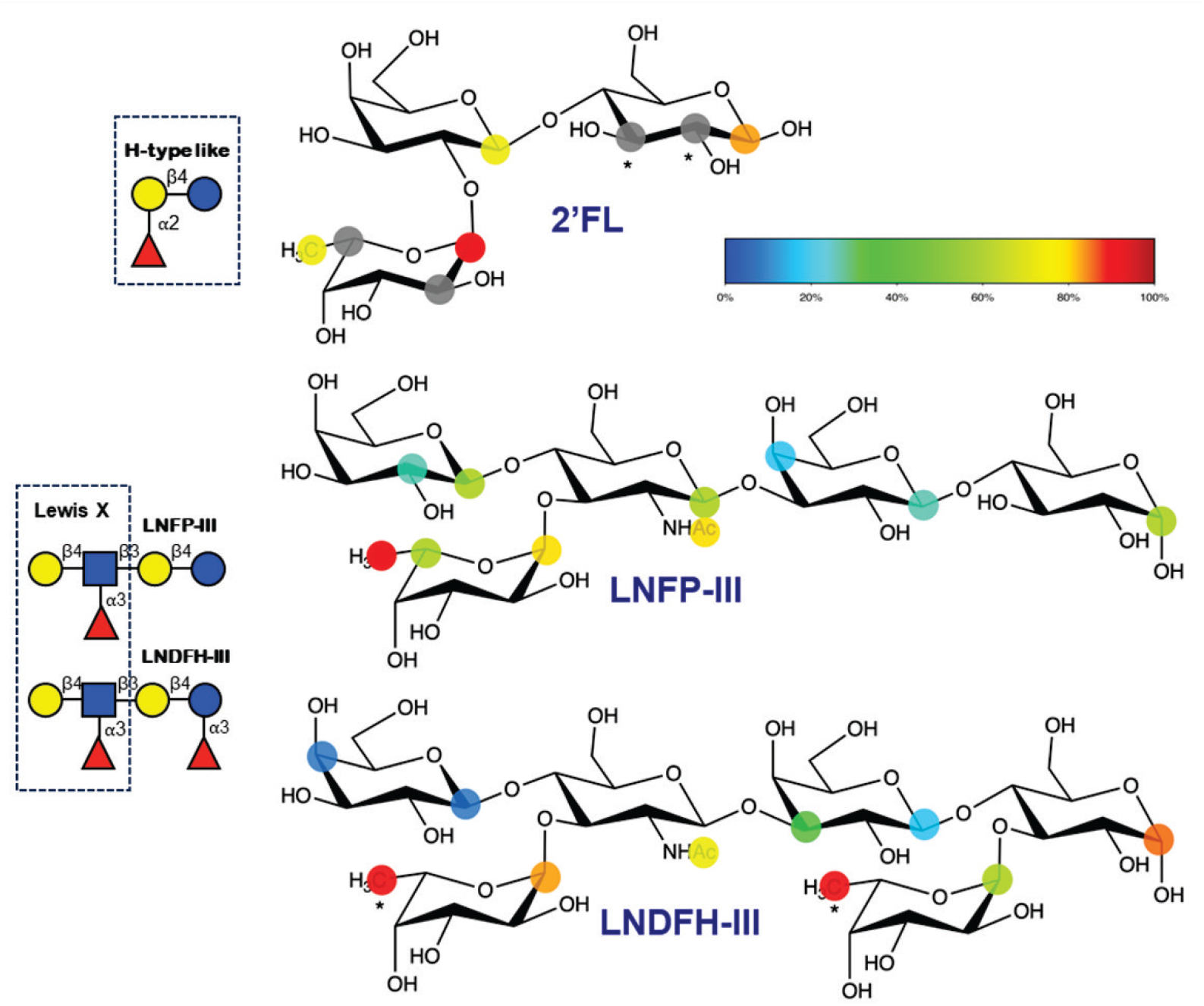
STD NMR binding epitope maps of the HMOs 2’FL, LNFP-III and LNDFH-III, to HuNoV GII.4 P-dimer. rd-STD NMR approach was used with t_sat_ = 0.75 and 6 s. The color codes represent the relative STD factors (normalized against the highest one), expressed as percentages of the individual ligand protons binding to P-dimer. Grey circles represent the interactions that were only observed at t_sat_ = 6 s, while asterisks indicate overlapping signals. Initial growth rates are shown in Table S1.

The main hotspots of the interactions for all three ligands were located at the Fuc residue, in particular at H1 in 2’FL and the methyl protons in LNFP-III and LNDFH-III (**Fig. 4**). For 2’FL, additional quantifiable contacts were visible in both H1^Glc^ and H1^Gal^ whereas unquantifiable contacts were visible in H2^Glc^ and H2^Fuc^, H3^Glc^ and H3^Gal^, and H5^Fuc^ due to significant signal overlap. Overall, all three residues of 2’FL contribute to binding to GII.4. Regarding LNFP-III binding to GII.4, each sugar residues established quantifiable interactions with the protein, supporting that longer HMOs can be accommodated into its glycan binding site. Noticeably, the lactose core showed reduced interaction compared to the Lewis X motif of the LNDFH-III. Since both H1α^Glc^ and H1β^Glc^ appeared well separated from other proton signals, analysis of both α and β anomers was feasible for LNFP-III and LNDFH-III. The binding of the α anomer of LNFP-III is likely preferred over the β one as H1α^Glc^ showed about 5-fold stronger STD intensity than H1β^Glc^. This α preference was also observed for LNDFH-III, in this case with a 4-fold stronger STD intensity for H1α^Glc^. The stronger relative STD value of H1α^Glc^ of LNDFH-III binding epitope compared to LNFP-III suggests that the additional Fuc residue at the reducing end enhances binding of the Glc residue **(Fig. 4)**. Moreover, as the signals of methyl protons of the reducing and non-reducing fucose residues overlap, they were treated as equivalent since fucose is the key structural element of binding and, hence, it can be sensibly assumed that both fucose residues interact with the P-dimer. Interestingly, because H1 protons of both fucoses were well resolved, we were able to conclude a weaker contact of the reducing fucose ring, due to their differences in relative STD values in the binding epitope map **(Fig. 4)**.

In summary, our data indicate that GII.4 P-dimer binds preferentially to HMOs presenting motif that mimics either the H-type antigen (2’FL) or the Lewis X antigen (LNFP-III and LNDFH-III). A preference for the α-configuration of the reducing end glucose is observed for the longer HMOs LNFP-III and LNDFH-III. These results highlight the binding promiscuity of the GII.4 Sydney variant toward a range of extended glycan epitopes beyond the fucose residue.

### Crystal structures of the P domain–HMO complexes

Co-crystallization of the GII.2, GII.4 and GII.17 P domains with the HMOs library yielded seven complex structures with GII.4 and GII.17 in which the ligand was unambiguously modelled at the conserved HBGA-binding interface located at the apical surface of the P2 subdomain dimer (11, 15, 29). These results are summarized in **Table 1** along with all the data collected on the different co-crystals obtained and are detailed thereafter.

**Table 1.**
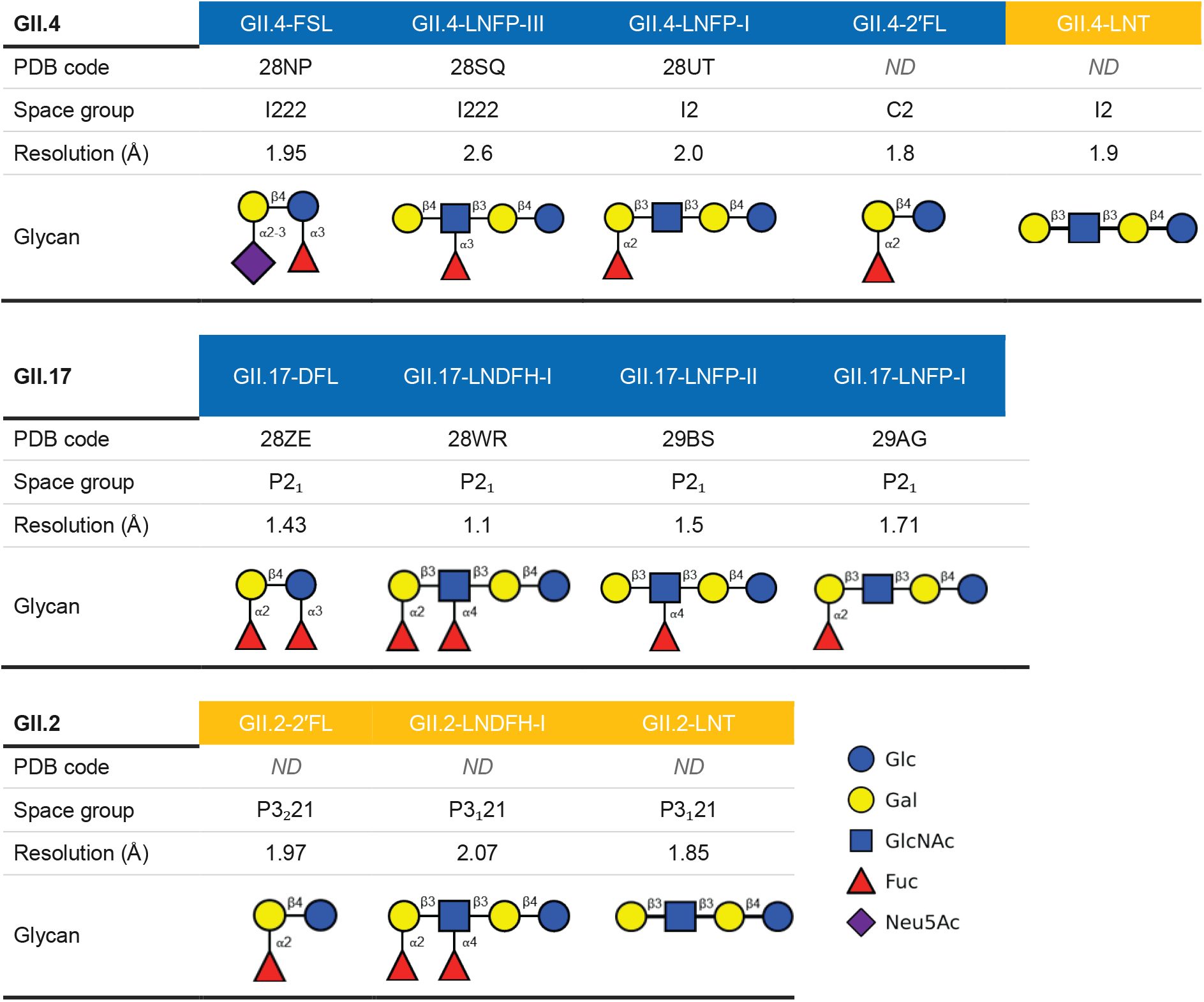
Summary on the data collected after co-crystallization screening of GII.4, GII.17 and GII.2 P domain with human milk oligosaccharides. The structures with the HMO bound were deposited to the PDB (blue header) contrary to those where the HMO did not bind corresponding to known apo structure (orange header). For GII.4-2′FL, the structure was not deposited as it already exists under code 8Y5V (30). Monosaccharide symbols follow SNFG convention. ND: not deposited.

| GII.4 | GII.4-FSL | GII.4-LNFP-III | GII.4-LNFP-I | GII.4-2′FL | GII.4-LNT |
|---|---|---|---|---|---|
| PDB code | 28NP | 28SQ | 28UT | *ND* | *ND* |
| Space group | I222 | I222 | I2 | C2 | I2 |
| Resolution (Å) | 1.95 | 2.6 | 2.0 | 1.8 | 1.9 |
| Glycan |  |  |  |  |  |
| GII.17 | GII.17-DFL | GII.17-LNDFH-I | GII.17-LNFP-II | GII.17-LNFP-I |  |
| PDB code | 28ZE | 28WR | 29BS | 29AG |  |
| Space group | P2<sub>1</sub> | P2<sub>1</sub> | P2<sub>1</sub> | P2<sub>1</sub> |  |
| Resolution (Å) | 1.43 | 1.1 | 1.5 | 1.71 |  |
| Glycan |  |  |  |  |  |
| GII.2 | GII.2-2′FL | GII.2-LNDFH-I | GII.2-LNT |  |  |
| PDB code | *ND* | *ND* | *ND* |  |  |
| Space group | P3<sub>2</sub>21 | P3<sub>1</sub>21 | P3<sub>1</sub>21 |  |  |
| Resolution (Å) | 1.97 | 2.07 | 1.85 |  |  |
| Glycan |  |  |  |  |  |

Glc
Gal
GlcNAc
Fuc
Neu5Ac

For GII.4, crystal structures were solved in complex with FSL, LNFP-I and LNFP-III at 1.95, 2.0 and 2.6 Å resolution, respectively (data collection and refinement statistics in **Table S2)**. Structure in complex with 2′FL was also obtained but not refined as it was consistent with previously published structures (30) and was used solely as an internal control. Diffraction data were also collected after cocrystallisation with DFL, LNFP-II and LNDFH-III but they were too low resolution or quality to be usable or solve the structure. After inspection of the electron density for GII.4 cocrystallised with non-fucosylated LNT, no binding was observed which underscored the importance of fucose for HMOs recognition by the norovirus P domain. This agrees with the results from the STD NMR screening of the library of HMOs.

For GII.17, crystal structures in complex with LNDFH-I, DFL, LNFP-II and LNFP-I were solved in P2₁ space groups at resolution between 1.10 and 1.71 Å resolution providing an exceptionally detailed view of the binding interactions at high resolution (**Table 1** and data collection and refinement statistics in **Table S2**). For GII.2, well-diffracting crystals were obtained with multiple HMOs including 2′-fucosyllactose, but the canonical binding site remained empty in all structures, consistent with structural constraints on HBGA recognition previously identified for this genotype (**Table 1**) (19, 20). No co-crystals were obtained for LNFP-V and LNDFH-III for any variant. With the exception of GII.4-LNFP-I, which contains two P domain homodimers per asymmetric unit, all six productive complexes crystallized with a single homodimer in the asymmetric unit. In all cases, unambiguous 2Fo-Fc and Fo-Fc electron density was observed for the fucose-containing portions of the HMO ligands at all symmetry-independent binding sites, allowing confident modelling of the bound conformations (**Fig. S2-S3**).

### Structural analysis of GII.4 P domain interactions with HMOs

Across all GII.4 complexed structures, the interactions with the fucose are conserved. In details, Arg345 donates sidechain guanidinium hydrogen bonds to O3 and O4. Asp374 engages O2 and O3 through bidentate carboxylate hydrogen bonding, Thr344 main chain oxygen makes a direct hydrogen bond with O4 and Gly443* (* from the neighboring protomer of the P-dimer) main chain nitrogen with the fucose ring oxygen. Tyr444* aromatic ring forms a CH-π stacking interaction with the fucose methyl group. Together, these five residues (Arg345, Asp374, Thr344, Gly443 and Tyr444) define the invariant recognition scaffold for fucose previously described for GII.4 across multiple epidemic variants and HBGA types (**Fig. 5A**) (17, 25, 31). This scaffold is further stabilized by a conserved water molecule that bridges the Fuc-O2 to the main chain nitrogen of Gly346 and the main chain oxygen of Cys441*. This bridge is resolved in the FSL and LNFP-I complexes but it is not modelled in the LNFP-III complex due to lower resolution and reduced solvent content that make an unresolved bridge more likely than a genuine absence. The interactions with the non fucose moieties of each HMO are as follow.

**Figure 5.**
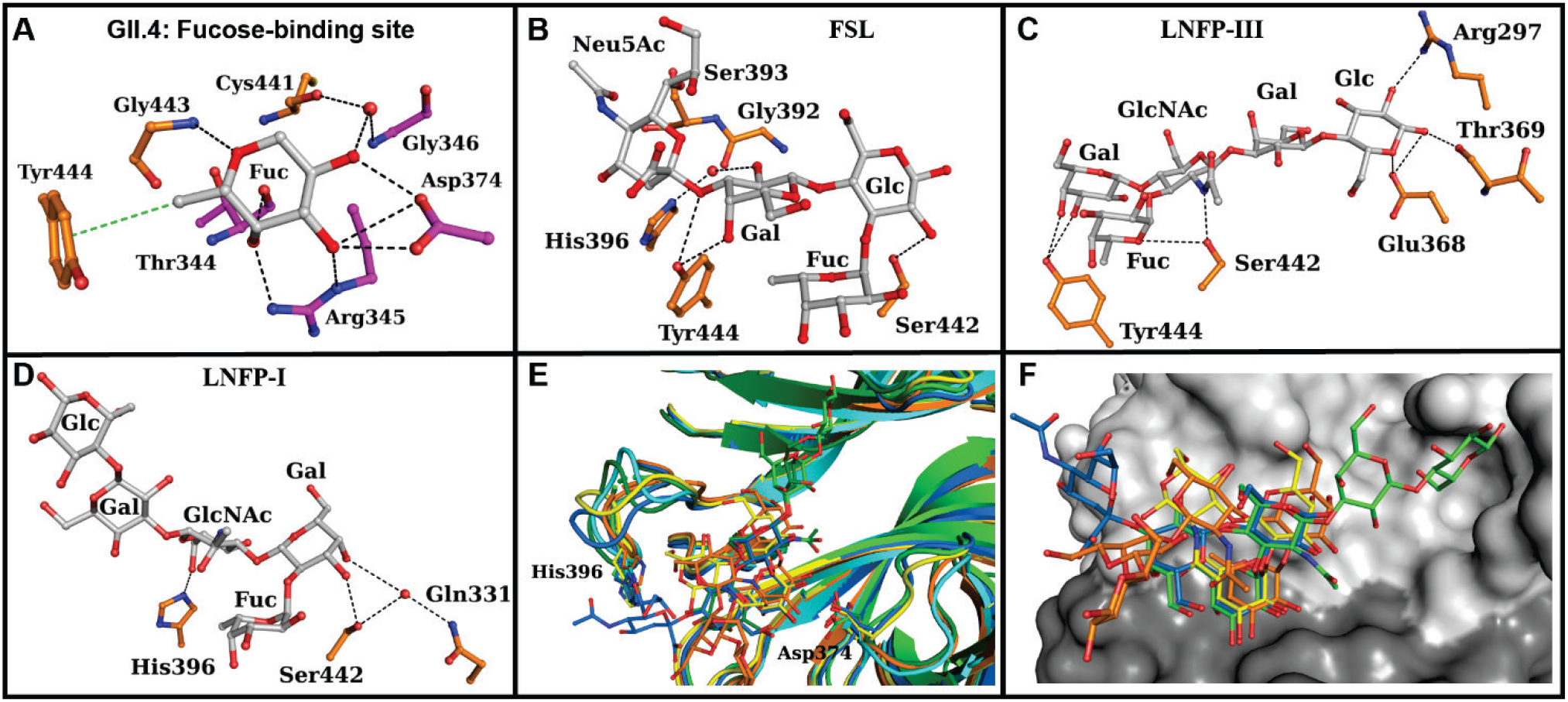
Structural basis of HMO recognition by the GII.4 P domain. (**A**) Zoom in the interactions in the fucose-binding site, showing its invariant recognition scaffold (**B–D**) Zoom in the interactions network for each HMO beyond the fucose moiety: (**B**) FSL (PDB 28NP), (**C**) LNFP-III (PDB 28SQ) and (**D**) LNFP-I (PDB 28UT). The protein carbons are colored according to the protomer in the dimer (orange or magenta) and water molecules are represented as red spheres. Hydrogen bonds and CH–π interactions are shown as black and green dashed lines, respectively. (**E-F**) Overlay of the FSL (blue and cyan), LNFP-III (green and dark green), LNFP-I (full pentasaccharide, orange and trisaccharide, yellow) complexes displaying surface loop movements (**E**) and binding surface (**F**). Ligands are displayed in ball and sticks and with their carbons colored as described. The surface is represented in different shade of grey according to protomer for LNFP-III.

In the FSL complex, a single binding site of the dimer is occupied. Tyr444* hydroxyl makes direct hydrogen bonds with Gal-O3 and Gal-O4, while His396* side chain interacts with Gal-O2 via a water bridge. The Glc-O2 forms a direct hydrogen bond with Ser442 hydroxyl and an additional contact with the fucose-bridging water described above (**Fig. 5B**). The sialic acid moiety only makes hydrophobic packing with Gly392*, Ser393* and His396*. No direct polar contacts to the sialic acid carboxylate group were observed and the interactions described are van der Waals, consistent with the peripheral positioning of this moiety relative to the core fucose-binding site (**Fig. 5B**).

In the LNFP-III complex, both binding sites are occupied and present equivalent interactions: the Fuc-O2 and GlcNAc amine are hydrogen bonded with Ser442* hydroxyl and the Gal-O3 with Tyr444* hydroxyl and Gal-O4 in one site (**Fig. 5C**). At one site, the glucose additionally forms direct hydrogen bonds with Arg297* (O2), Glu368* (O1) and Thr369* (O1). The electron density for this terminal glucose is, however, less well-defined than for the preceding moieties, likely reflecting increased conformational flexibility at the terminus of the sugar chain. These contacts should therefore be interpreted with caution, but it is noticeable that interaction with the glucose was also seen in the STD NMR experiment (**Fig. 4**). It is to be noted that the side chain of Arg297* rotates around CD-CG bound to avoid clashes with the glucose moiety and optimize hydrogen bonding.

In the LNFP-I complex, three of the four binding sites are occupied across the two dimers in the asymmetric unit, with the fourth site showing no interpretable electron density for the ligand that was therefore left unoccupied in the final model. The full pentasaccharide could only be modelled in one protomer where the lactose moiety is stabilized by crystal contacts that will not be further detailed as they may be not representative. In the other protomers, the lactose moiety is fully solvent exposed and could not be modelled. The Gal-O3 consistently contacts Ser442* side chain and a variable additional water bridge can be seen between Gal-O4 and the main chain oxygen of Gln390*. The GlcNAc-O6 is hydrogen bonded with either the Gln390* side chain oxygen or the His396* imidazole ring nitrogen (**Fig. 5D**).

Superposition of the three GII.4 complexes shows that the surface loops containing Asp374 (residues 367–381) is displaced in one protomer but it seems due to crystal contacts rather than ligand binding. The others protomers adopt the same conformation seen in FSL and LNFP-III complexes. There is however a rigid body translation of about 0.7 Å of the neighboring protomer leading to the same shift in the sugar when the full ligand is modelled (**Fig. 5E-F**). Despite this displacement, all interactions with the fucose are retained showing how small accommodations can be made. Asp374 remains hydrogen bonded to the fucose ring in all three complexes despite different orientation, though the bidentate contact to both O2 and O3 seen in FSL and LNFP-III narrows to a single close contact in LNFP-I. The beginning of the loop carrying His396* (residues 389*-399*) can present different conformations and in some protomers could not be modelled (**Fig. 5E**). The His396 imidazole lies 3.1–3.5 Å from the bound glycan in the FSL and LNFP-I complexes, compatible with the direct and water-mediated contacts described above, but 5.5–6.2 Å away in both LNFP-III binding sites, where this region makes no contact with the ligand. When FSL is bound, it also adopts a different confirmation to optimize hydrophobic interactions with the sialic acid moiety notably with Gly392* (**Fig. 5B and 5E**). The different ligands can bind to a broad surface area around the fucose finding site that cover six additional sugar moieties and whose shape is dictated by the conformation of those surface loops (**Fig. 5F and S2**).

### Structural analysis of GII.17 P domain interactions with HMOs

Across the four GII.17 complexed structures, both binding sites are occupied. The fucose recognition mirrors the scaffold identified for GII.4, involving the following conserved amino acids: Arg349, Asp377, Thr348, Gly441 and Tyr442 **(Fig. 6A)**. This scaffold is further stabilized by a two-waters bridge to the Fuc-O2. The first water is jointly coordinated by the main chain nitrogen of Ser350 and the main chain oxygen of Ser439*, a pairing conserved across all four GII.17 structures. This water is linked to a second water that contacts the Fuc-O2 directly. In contrast to GII.4 where Tyr444 engages downstream sugars through direct hydrogen bonds, Tyr442 in GII.17 participates exclusively through water-mediated contacts with the galactose and GlcNAc moieties. A further GII.17-specific feature is Gln379, which is water bridged with Fuc-O3 and to downstream sugar moieties across all the four structures with no equivalent in GII.4. The per-complex additional contacts are described below.

**Figure 6.**
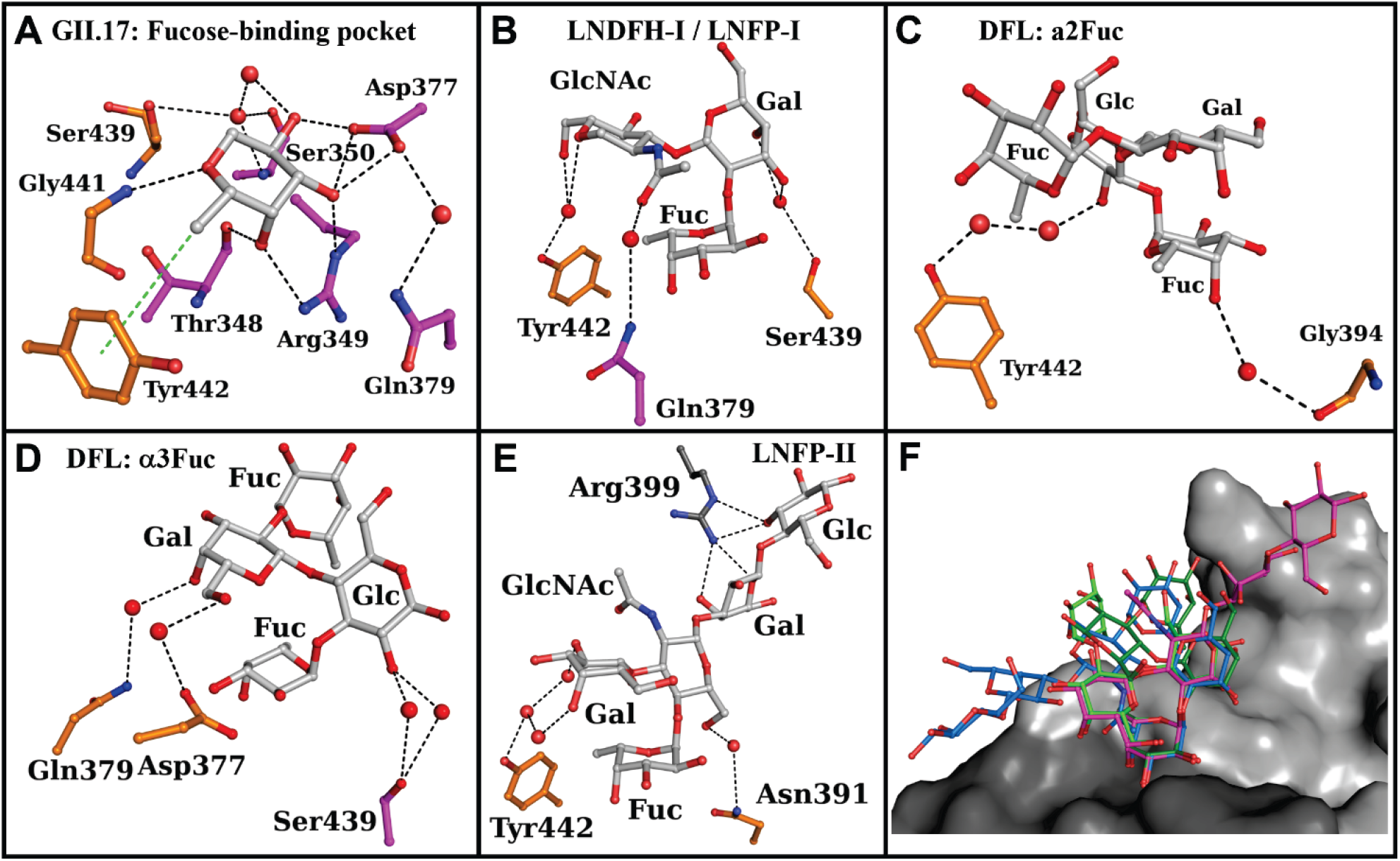
Structural basis of HMO recognition by the GII.17 P domain. **(A)** Zoom in the interactions in the canonical fucose binding site. **(B, D–F)** Interaction network for each HMO beyond the fucose-binding scaffold: **(B)** LNDFH-I and LNFP-I (PDB 28WR and 29AG, respectively), **(C)** DFL (PDB 28ZE) with Fucα1-2Gal as anchor, **(D)** DFL with Fucα1-3Glc as anchor and **(E)** LNFP-II (PDB 29BS). Terminal monosaccharide moieties that make no interactions with the protein were omitted for clarity in **B**. The protein carbons are colored according to the protomer in the dimer (orange or magenta) and water molecules are represented as red spheres. Hydrogen bonds and CH–π interactions are shown as black and green dashed lines, respectively. **(F)** Superposition of LNDFH-I (α2Fuc, blue), DFL (α3Fuc, green) and LNFP-II (magenta, α4Fuc) on GII.17 P domain surface colored by protomer in different grey shades.

In LNFP-I and LNDFH-I complexes, the α2Fuc is recognized and water mediated interactions are only observed for the Gal-GlcNAc disaccharide as follow. Both GlcNAc-O5 and GlcNAc-O6 contacts Tyr442* hydroxyl through the same water similarly to Gal-O3 and Gal-O4 with the Ser439* main chain oxygen. Gln379 side chain nitrogen interacts with the GlcNAc acetyl oxygen through one or two water molecules depending on the binding site **(Fig. 6B)**. The lactose moiety does not interact with the P-dimer and is exposed to the solvent. In consequence, the glucose moiety was too disordered to be modelled in the LNFP-I structure. The additional α4Fuc does not displays any interactions but adopts a close conformation with the galactose known to be a low energy conformation and brings more rigidity to the whole hexasaccharide (32). This seems specially to stabilize the lactose moiety which becomes more ordered and could be modelled in the electron density in LNDFH-I and not LNFP-I. Overlay of both oligosaccharides shows similar overall configuration but slight rotation up to 15° rotation around the O1’-C2 bond of the α(1–2) glycosidic linkage can be observed bringing the remaining oligosaccharide closer to the protein surface and placing the GlcNAc ring perpendicular to the Tyr442 phenyl ring. Superposition of the α1-2Fuc-anchored LNDFH-I and LNFP-I complexes shows that both the protein scaffold and the bound sugars are essentially superimposable on the shared fucose-galactose anchor, with no accompanying change in the protein (**Fig. S4**).

In the DFL complex, both binding orientations of this difucosylated HMO are captured within the same crystal, making this the only complex where the two binding sites differ meaningfully and warrant separate description. When DFL is anchored via the α2Fuc, the α3Fuc-O4 makes a water bridge with the Gly394* main chain oxygen and Glc-O2 with Tyr442* hydroxyl **(Fig. 6C**). The ligand path diverges beyond the galactose, where DFL continues onto a different sugar (Glc instead of GlcNAC) via a different linkage than LNFP-I and LNDFH-I with slight rigid body on surface loop with no impact on sugar interactions (**Fig. S4B**). The only significant difference in the binding site is the rigid-body movement of Tyr442 side chain of 0.8 Å to accommodate the binding of the additional αfucose in LNDFH-I and DFL **(Fig. S4B)**. When the α3Fuc occupied the fucose binding site, Gal-O4 and Gal-O6 are water-bridged to the Gln379 and Asp377 side chains, respectively and Glc-O2 is water-bridged to the Ser439* main chain oxygen **(Fig. 6D)**. The α2Fuc is solvent exposed and doesn’t make additional interactions.

In the LNFP-II complex, the full pentasaccharide could be modelled in one site whilst only the Lewis a trisaccharide in the other. The proximal galactose O2, O3 and O4 are water bridged with Tyr442* hydroxyl through one or two water molecules. GlcNAc-O6 presents a weak hydrogen bond with the amide nitrogen of Asn391 (3.5 - 3.75 Å). Glc-O3 is hydrogen bound with Arg399 guanidinium from a symmetric molecule stabilizing it resulting in good electron density for the modelling of the lactose moiety **(Fig. 6E)**. Subtle changes are observed with in particular in the loop comprising Gly394-Pro397 with a rigid-body movement of 1.5 Å of Asn395 main chain and rotation of almost 90° of its side chain away from the binding site in order to accommodate the lactose moiety. GII.17 P domain bind the HMOs indifferently of the fucose linkage and the ligand span a very extensive binding surface with seven additional subsites that is highly solvent exposed **(Fig. 6F)**.

### Comparative structural analysis

Despite sharing only approximately 60% sequence identity across the P domain, GII.4 and GII.17 employ a structurally conserved recognition scaffold for fucose involving five conserved residues as mentioned above **(Fig. 7A)**. The geometry of the CH-π contact with the conserved tyrosine differs subtly but consistently between the two genotypes. In GII.4 the C6 methyl-to-ring centroid distance ranges from 4.0 to 4.4 Å, comfortably within the canonical range for CH-π interactions (33, 34), whereas in GII.17 the equivalent distances are larger and range from 4.4 to 4.7 Å **(Fig. 7B)**. Whether that translates into any difference in the enthalpic contribution of this contact cannot be assessed from the present structural data alone. Beyond the CH-π contact, the two tyrosine residues diverge further in their engagement with downstream sugar moieties. In GII.4, the tyrosine hydroxyl is involved in a direct hydrogen bond with the galactose O3 and O4 of FSL and LNFP-III **(Fig. 5B-5C)** whilst in GII.17, it participates exclusively in water-mediated contacts with the GlcNAc and/or the galactose moieties of LNFP-I, LNDFH-I and LNFP-II **(Fig. 6B, 6E)**.

**Figure 7.**
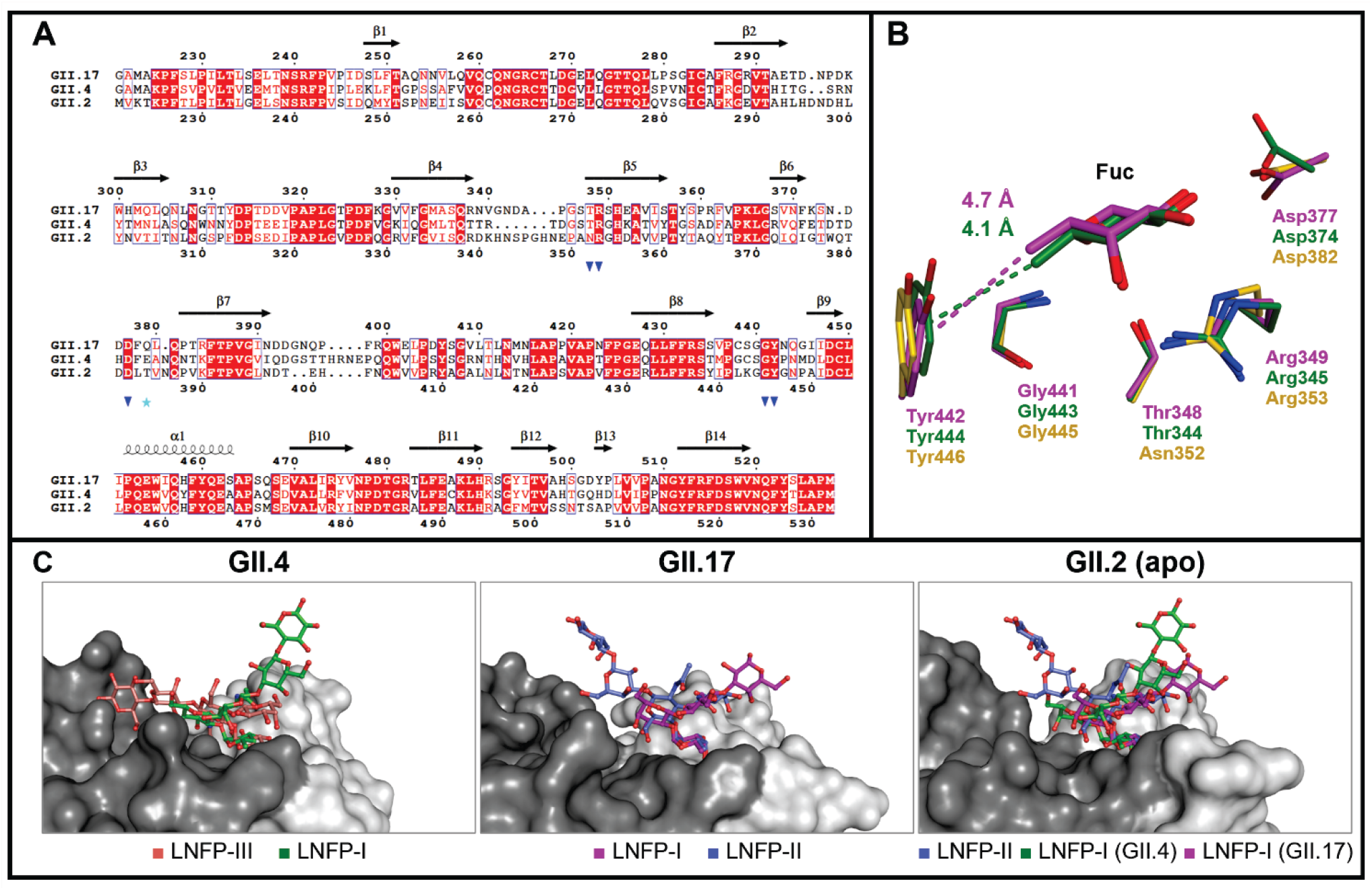
Comparative structural analysis of the fucose-binding scaffold in GII.4, GII.17 and GII.2 P domains. **(A)** Structure-based sequence alignment of the GII.17, GII.4 and GII.2 P domains, generated with ESPript 3.0 (35). Secondary structure elements derived from the GII.17 structure are displayed above the alignment. Blue triangles indicate residues conserved in fucose recognition and the cyan star denotes Gln379, involved in a conserved water-mediated bridge in the GII.17. **(B)** Superposition of the GII.4 (green), GII.17 (magenta) and GII.2 apo (yellow) fucose-binding site after pair-fit alignment on five conserved residues, showing the CH–π stacking interaction (magenta and green dashed lines). **(C)** Surface representation of the fucose-binding site in GII.4, GII.17 and GII.2, illustrating differences in binding-site architecture and solvent exposure across the three genotypes; the two protomers of each dimer are shown in two shades of gray. Left, GII.4 superposed with LNFP-III (pink) and LNFP-I (green). Middle, GII.17 superposed with LNFP-I (magenta) and LNFP-II (blue). Right, apo GII.2, equivalent to the previously published structure (PDB 5YSX), with LNFP-II (blue) together with LNFP-I from the GII.4 (green) and GII.17 (magenta) complexes.

The overall architecture of the P-dimer binding surface differs however, between the three genotypes as the surface loops enclosing it (β4-β5, β5-β6, β7-β8 and β8-β9) which present divergence in sequence, length and conformation **(Fig. 7A-C** and **S5)**. The β4-β5 loop carries a three-residue insertion in GII.17 relative to GII.4 (339–347 versus 338–343) that enlarges the contact area around the fucose subsite, while the β6-β7 loop anchors Asp374 in GII.4 and the equivalent Asp377 in GII.17; within this loop, Gln379 in GII.17 occupies the position spatially equivalent to Glu376 in GII.4. This creates divergent secondary contact surface that impacts on the fine specificity for their ligand. The whole binding site is shaped as in a groove that is less extended at the reducing end for GII.2 whilst GII.17 is shallower and more solvent exposed and GII.4 is more extended and enclosed **(Fig. 7C)**. This allows all genotypes to bind a broad spectrum of fucosylated ligands without preference for a linkage. As a consequence, the crystal structures shows that GII.4 relies on direct sidechain contacts (Tyr444*, Ser442*, Gln390*, His396*) for secondary recognition whereas GII.17 achieves equivalent coverage primarily through an ordered solvent network anchored by Gln379, Ser439 and Tyr442* that appears to substitute structurally for the direct sidechain contacts seen in GII.4. Within the β7-β8 loop that carries Gln390 and His396 in GII.4, this region closes over the bound glycan in the FSL and LNFP-I complexes, positioning the His396 imidazole 3.1–3.5 Å from the ligand, but remains disengaged in the LNFP-III complex (5.5–6.2 Å). The equivalent GII.17 loop, carrying Asn391 and Gly394-Pro397 reorganizes around the lactose moiety of LNFP-II as described above.

LNFP-I, present in the crystallization panel of both GII.4 and GII.17, allows a direct comparison on an identical ligand. In GII.4, His396 (β7-β8 loop) side chain adopts a different conformation to improve binding through a hydrogen bond with the GlcNac-O6 when the lactose moiety is ordered. In GII.17, the equivalent residues (Asn391, Gly394-Pro397, same loop) remain disengaged at both occupied sites (6.5 Å) regardless of how much of the ligand is ordered. The glucose moiety itself is modelled in one of three GII.4 sites but in neither of the two GII.17 sites, consistent with the more rigid, solvent-mediated GII.17 surface accommodating the reducing end of LNFP-I less readily than GII.4’s **(Fig. S5)**. The most notable genotype-specific feature is the role of Gln379 in GII.17, which makes water bridges with both fucose O3 and downstream GlcNAc or galactose moieties across all the four complex structures **(Fig. 6B-F).** These results show that GII.4 and GII.17 recognise longer fucosylated HMOs through the same fucose site used for HBGAs, which supports the decoy hypothesis, whereas GII.2 did not bind them under our conditions despite an intact scaffold.

### Backbone NMR assignment of GII.4 Sydney P domain

The perdeuterated ²H/¹³C/¹⁵N-labelled GII.4/Sydney/2016 P domain produced a well-dispersed ¹H–¹⁵N TROSY-HSQC spectrum **(Fig. 8A)**, consistent with a folded protein. NMR experiments for backbone resonance assignments were performed using an TROSY-based pulse sequences due to the large molecular size of the dimeric P domain (69 kDa). To facilitate assignment, the spectra were analyzed by comparison with the published assignment of the closely related GII.4 Saga P domain (BMRB 27445) (36).

**Figure 8.**
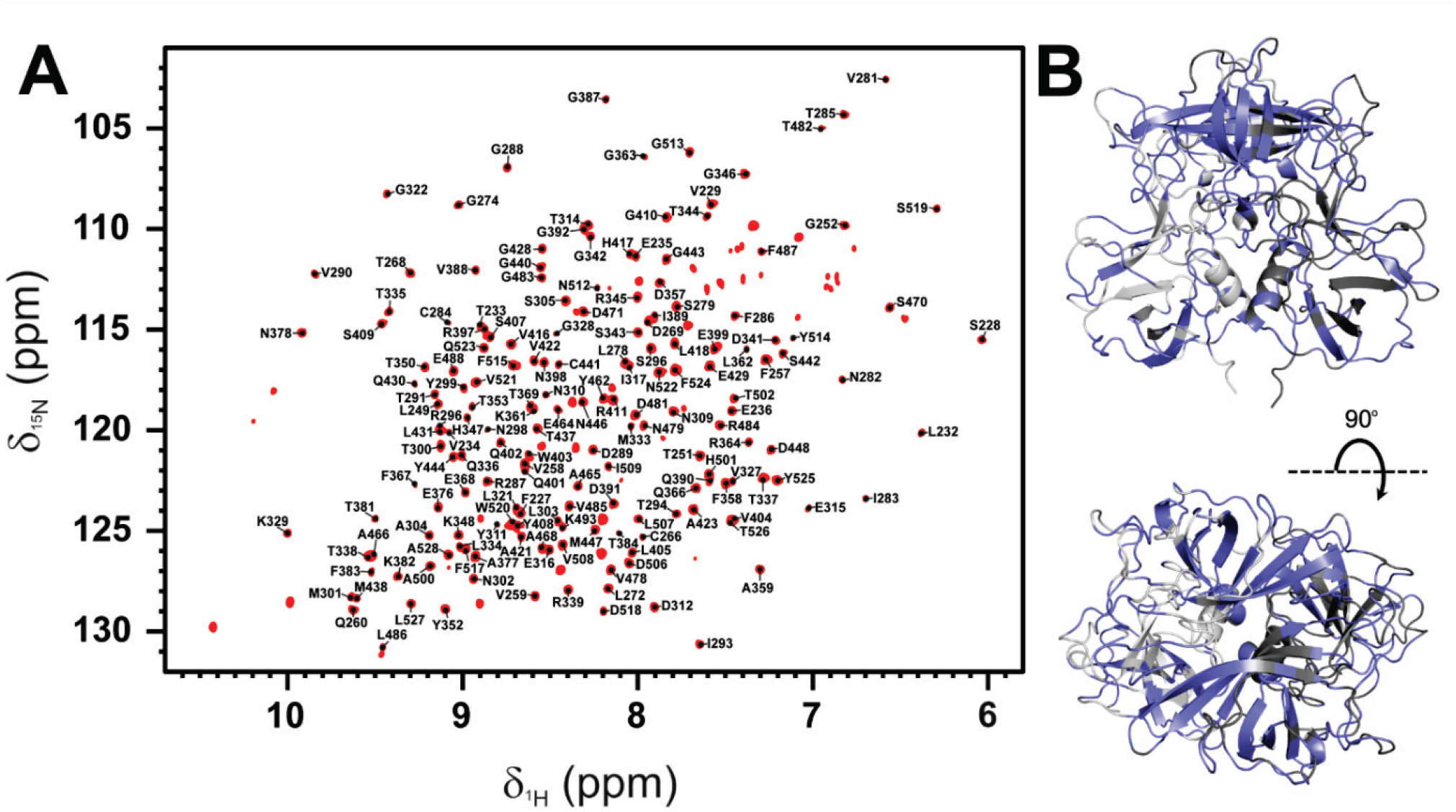
Backbone assignment of the GII.4 P domain. (A) 1H-15N TROSY-HSQC spectrum of the 2H/13C/15N enriched sample with assigned resonances. (B) Assignment coverage mapped onto the apo P domain dimer structure, shown in top and side views. Assigned residues are highlighted in blue. The overall completeness reached approximately 62%, demonstrating substantial coverage of the canonical glycan-binding site.

This combined analysis enabled the assignment of approximately 62% of the backbone amide resonances; those remaining resonances were unassigned due to signal overlap, low intensity or severe line broadening. Importantly, most residues located in the canonical glycan-binding site were reliably assigned **(Fig. 8B)**. Therefore, although the overall assignment coverage was incomplete, the assigned resonances were sufficient to support residue-specific chemical shift perturbation (CSP) analysis of ligand binding.

### CSPs and structural mapping of the binding interface

To investigate glycan recognition by GII.4 Sydney P domain, ¹H–¹⁵N TROSY-HSQC titration experiments were performed using the selected HMOs (2′FL, 3FL, FSL and LNFP-I). Most resonances exhibited no significant changes, indicating that the protein retains its structural integrity upon ligand binding. All the investigated ligands caused continuous cross-peaks shifts of the affected residues during the titration, characteristic of the fast exchange regime on the NMR timescale. CSP analysis revealed both shared and ligand-specific perturbation patterns **(Fig. 9A)**.

**Figure 9.**
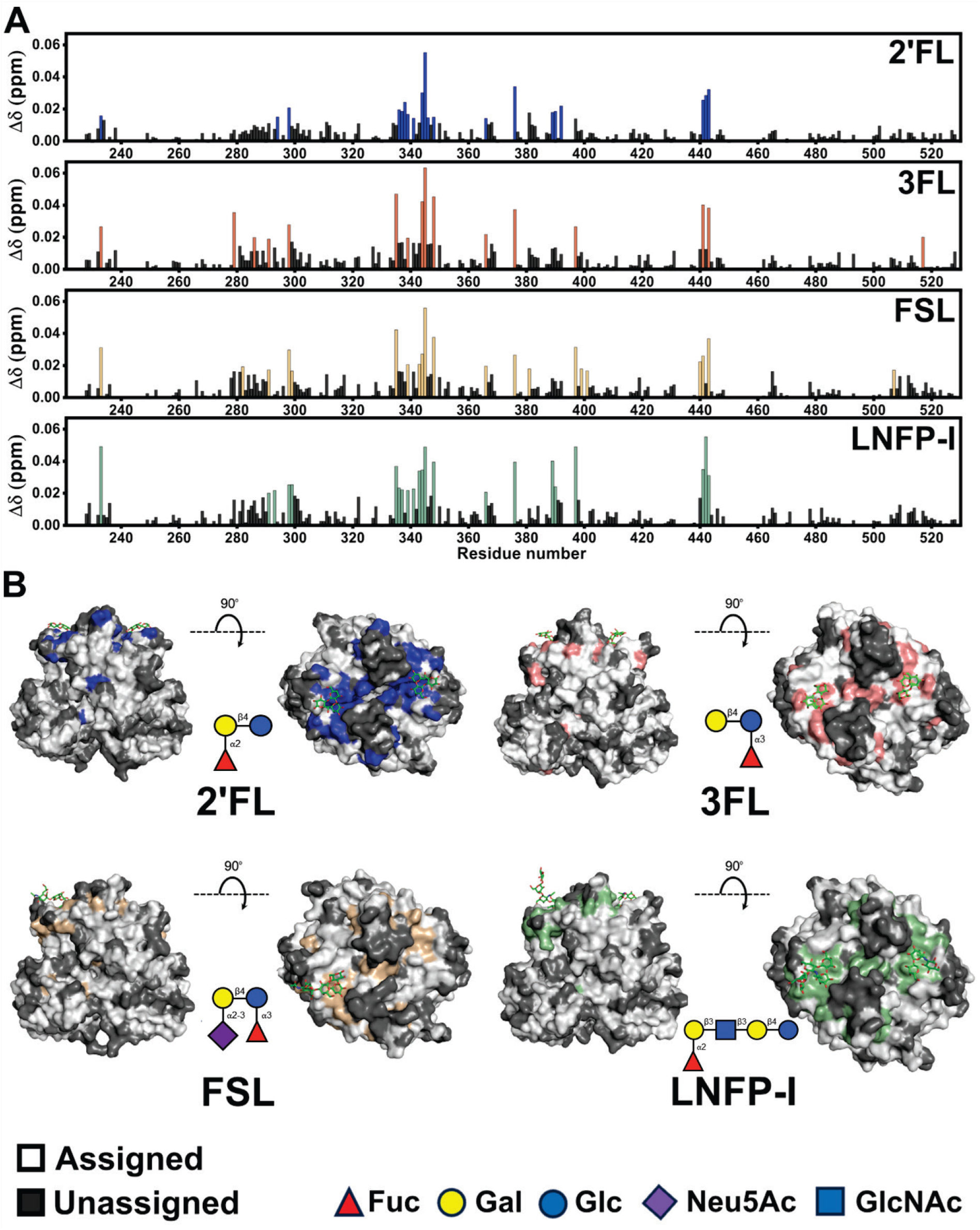
(A) Chemical shift perturbation (CSP) profiles of the norovirus GII.4 P domain dimer induced by 2’FL (blue), 3-FL (red), FSL (pale orange), and LNFP-I (green) with each ligand evaluated at a concentration of 16 mM. (B) Mapping of the perturbed residues at the concentration of 16 mM onto the crystal structures of the P domain (color scheme as in A, 2’FL (PDB 8Y5V), 3FL (PDB 5IYW), FSL (PDB 28NP) and LNFP-I (PDB 28UT).

In general, the highest chemical shift perturbations clustered in the top loop and β-sheet region of the P-dimer. The affected residues included Thr344, Arg345, Ser442 and Gly443, consistent with the known glycan-binding site. In addition, secondary CSPs were observed at distal residues, suggesting possible allosteric effects within the domain due to ligand binding. Similar effects have also been observed in the GII.4 Saga P domain (37). Mapping the significantly perturbed residues onto the corresponding crystal structure of the complexes revealed a clear clustering around the surface-exposed loops that form the canonical glycan-binding interface **(Fig. 9B)**. Although all ligands engaged the protein in the same general binding region, FSL and LNFP-I induced more extensive perturbations that extended beyond the core fucose binding site **(Fig. S5)**.

Global fitting yielded dissociation constants in the millimolar range **(Fig. 10)**, consistent with the weak protein–glycan interactions typically reported for norovirus capsid proteins (38). Among the tested ligands, 3-FL showed the slightly stronger binding affinity (K_D_ ≈ 20 mM), followed by LNFP-I (K_D_ ≈ 22 mM) and 2′FL (K_D_ ≈ 24 mM), whereas FSL exhibited weaker binding (K_D_ ≈ 33 mM). Titrations for 2′FL and 3-FL were extended up to 128 mM, while those for FSL and LNFP-I were limited to 64 mM due to increasing resonance overlap between ligand and protein signals in the TROSY-HSQC spectra at higher concentrations.

**Figure 10.**
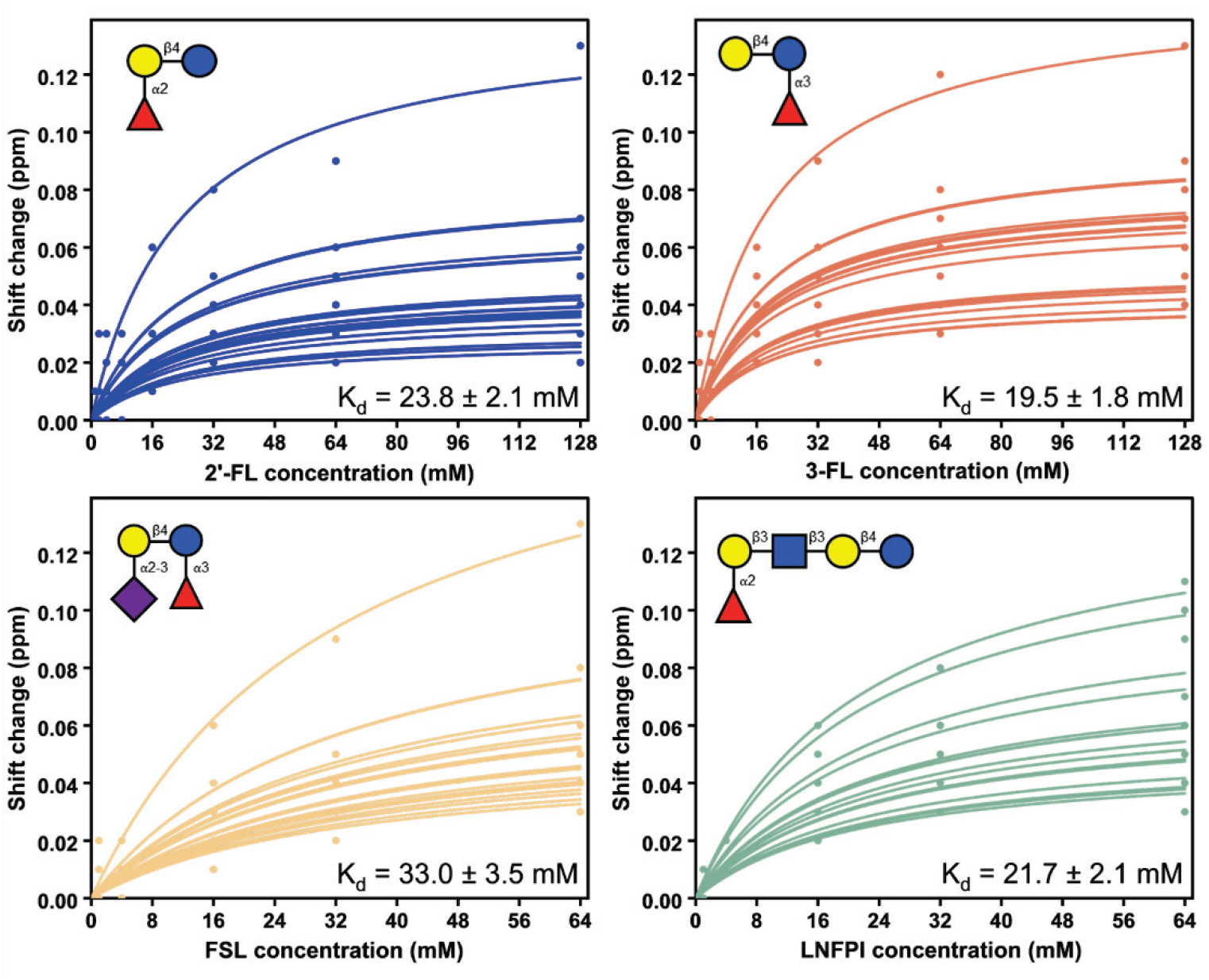
CSP titration curves and corresponding fitted used to determine the dissociation constants (K_D_) of 2’FL (blue), 3-FL (red), FSL (orange) and LNFP-I (green) for norovirus GII.4 P domain. The detailed fitting is shown in **Fig. S6**.

## DISCUSSION

This study provides the first high-resolution view of how the norovirus P domain binds naturally occurring HMOs across three antigenically and phylogenetically distinct GII genotypes. It extends previous structural studies on the trisaccharides 2′FL and 3FL and HBGAs, with HMOs comprising four to six monosaccharide units, two difucosylated and one sialylated species (25, 26). Here, the seven crystal structures described show anchoring though one fucose unit by the same five invariant amino acids scaffold as originally defined for HBGA trisaccharides (15–17). The fucosylated ligand can elongate both on the non-reducing and reducing ends of the core fucose thanks to a secondary binding surface that can also accommodate long and branched HMO like the difucosylated hexasaccharide LNDFH-I. α1-2, α1-3 and α1-4 linked fucoses were all found in the core fucose binding site and for the first time in a difucosylated ligand (DFL), both orientations were trapped and not only the α1-2 linked fucose as in GII.17 with LNFDH-I (28WR) or GII.4 with Lewis b (4OPO) (17). A large variety of fucosylated ligands can be tolerated as the binding surface is quite shallow and solvent-exposed. The STD NMR screening reaches the same conclusion from an independent, solution state perspective: every fucosylated HMO tested produced a detectable signal against GII.4 and GII.17, whereas the non-fucosylated ligands LNT and LNnT showed no significant signals. This confirms the crystallographic finding that engagement of the conserved binding site by fucose is obligatory rather than one of several alternative anchoring modes. Hydrogens bonds and CH–π stacking are limited to the fucosylated trisaccharide and are scarce for the Gal, Glc/GlcNAc whilst the lactose and second fucose moieties face the solvent and are found stabilized by crystal contacts **(Fig. 5-6)**. However, interactions with the reducing glucose were observed by STD-NMR **(Fig. 3)**.

We demonstrated that the surface loops enclosing the binding surface can adopt different conformations and adapt to the ligand nature to avoid steric clashes and optimize, when possible, interactions or hydrophobic contacts **(Fig. 5E**). They diverge between the genotypes and creates a different binding surface shaped as a groove that is more or less extended, shallow and solvent exposed **(Fig. 5E, 7C** and **S2-S3)**. These structural differences are echoed in the STD NMR data: 3FL and LNFP-V bind more strongly to GII.17, whereas LNFP-III, which presents the same fucosylated motif at the non-reducing end on a GlcNAc rather than at the reducing-end glucose, shows the opposite preference. Together, these results show that although GII.4 and GII.17 recognize fucose through an identical mechanism, the changes in binding surfaces rigidity and extent will define the fine specificity and selectivity as well as perhaps affinity for each genotype on a given HMO.

Beyond the shared fucose core, GII.4 and GII.17 diverge in the plasticity of the surface that reads out the remaining sugar units. GII.4 achieves secondary recognition through a loop capable of closing around the ligand in a sequence-dependent manner, engaging downstream sugars directly when ligand geometry permits. GII.17 instead relies on a fixed, ordered solvent network that does not reconfigure with the ligand. This contrast offers a structural rationale for the genotype-specific preferences captured independently by STD NMR, where the same fucosylated motif elicited opposite relative intensities in GII.4 and GII.17 depending on its position within the HMO. Whether the more rigid GII.17 interface also constrains binding kinetics or affinity, as opposed to specificity alone, cannot be resolved from the present structural data.

The weak and limited interactions as well as the shallow binding surface probably explain the weak affinity obtained by protein-based NMR titrations with GII.4 that also provided a quantitative and residue resolved layer of validation. It confirmed that the dominant solution state binding event corresponds to the interactions captured in the crystal structures in the core fucose binding site. In particular, Thr344, Arg345 and Gly443, these residues of the defined fucose-recognition scaffold by crystallography, were consistently perturbed by 2’FL, 3FL, FSL and LNFP-I (**Fig. S7**). Perturbations were also observed around the Asp374-containing loop, with residues Glu376 or Asn378 affected depending on the ligand, further supporting engagement of the same conserved binding region in solution. Crystallization did not seem to select an alternative or non-physiological glycan orientation. Meanwhile, the CSP profiles reveal differences in the spatial extent of the protein response. That FSL and LNFP-I perturb a broader surface beyond this core pocket than the shorter 2′FL and 3FL is consistent with a more extensive binding surface as resolved by X-ray crystallography for these longer ligands. For LNFP-I, perturbations at residues Ile389, Gln390, Arg397, Cys441-Gly443 are consistent with involvement of the secondary binding surface containing Gln390, His396 and Ser442. FSL similarly perturbed residues Arg397, Glu399 and Gln401, close to the His396-containing region involved in the peripheral packing of its extended glycan chain and sialic acid moiety. 2′FL also perturbed residues beyond the central fucose-recognition residues, consistent with the additional contacts involving the non-fucose portion of the ligand observed by STD NMR. And also, galactose and GlcNAc contacts common to both and the sialic acid packing unique to FSL (38).

However, a broader perturbation profile does not necessarily indicate stronger binding. Chemical shifts are sensitive not only to direct intermolecular contacts but also to indirect structural and dynamic changes. Consequently, an extended CSP footprint may arise from several weak secondary interactions or from ligand-induced redistribution of local conformational states without producing a proportionate gain in binding free energy (39). Global fitting of titration curves yielded dissociation constants from 20 to 33 mM placing HuNoV–HBGA interactions in the low millimolar range, critically correcting earlier affinity estimates that had been substantially overestimated from STD NMR competition assay (38). Notably, FSL does not bind more tightly than the simpler 2’FL or 3FL despite forming a wider network of secondary contacts, indicating that an expanded contact surface does not, at least for this subset of ligands, translate into a proportionate gain in affinity. This is consistent with the crystal structure of the FSL complex: compared to 3-FL, which shares the same Fucα1-3Glc linkage, FSL’s adds a sialic acid residue. This moiety engages the protein exclusively through hydrophobic packing with Ser393* and His396* rather than through direct polar contacts, offering a structural rationale for why the additional contact surface does not translate into higher affinity.

Attempts to measure these interactions independently by isothermal titration calorimetry (ITC) and by biolayer interferometry (BLI), with the P domain immobilized on the sensor chip for the latter, did not yield quantifiable binding data for either genotype (data not shown). For ITC, the absence of a measurable signal is more likely attributable to a small or unfavorable enthalpy change accompanying binding than to affinity alone, although a genuinely weak interaction cannot be excluded as a contributing factor. For BLI, the most likely explanation is that dissociation occurred too rapidly for the instrument to resolve an off-rate, precluding determination of a dissociation constant, though other causes of the failure to obtain a fit cannot be ruled out. Characterization of these interactions required overcoming the considerable challenges associated with solution NMR analysis of the 69 kDa P-dimer. Despite perdeuteration, high-field TROSY experiments and the acquisition of several triple-resonance datasets, only 62% of the backbone amide resonances could be assigned. Similar difficulties have been previously reported for the closely related GII.4 Saga P-dimer, for which short transverse relaxation times, incomplete amide proton-deuterium exchange and local conformational exchange affected spectral quality (37). Nevertheless, the assignment coverage obtained here included most residues surrounding the glycan-binding site. Consequently, the partial assignment was sufficient to identify the principal interaction surface and to monitor binding quantitatively.

In GII.2 P domain complexes with multiple HMOs, including 2′FL, the canonical HBGA binding site remained unoccupied despite the five-residue fucose-binding scaffold adopting the conformation observed in GII.4 and GII.17. This in contrast with the hypothesis that the lack of HBGA binding to GII.2 was related to a different conformation of an aspartate side chain away from the binding site as observed in the apo structure of the GII.2 Snow Mountain strain (20). Superimposition of the three genotypes shows that GII.2 has the least extended and more open binding surface **(Fig. 7C)** but whether this accounts for the absence of stable occupancy in GII.2, independently of the intact fucose recognition scaffold, remains an open question. The STD NMR data are consistent with the absence of stable occupancy at this site: almost every HMO produced signal at or below the limit of detection against GII.2, in clear contrast to the signals obtained for the same ligands against GII.4 and GII.17. The convergence of two independent methods on the same result argues against a crystallization artefact, but the present data cannot distinguish between an intrinsic too low affinity to support stable ligand occupancy under the conditions used here and a requirement for an additional factor absent from our experiments. Bile constituents such as bile salts have been shown to enhance HBGA engagement by other GII.2 strains despite capsid microvariation (9). Although not tested in the present study, this offers a plausible explanation for how the binding site remains unoccupied in absence of cofactors (9). This will be tested in the near future.

## CONCLUSION

These results extend the structural basis of norovirus recognition of HMOs beyond simple fucosylated trisaccharides to a structurally diverse panel of longer, naturally occurring HMOs. They demonstrate a broad binding profile of HMOs and help to interpret differences in inhibitory potency across the most common norovirus lineages examined. They further show that the proposed decoy-like mimicry of HBGAs by HMOs (22–24) relies on a fucose recognition mechanism that is conserved between GII.4 and GII.17, while being fine-tuned by genotype-specific secondary contacts. In contrast, GII.2 remains refractory to HMO binding in the set-up used despite displaying a functional fucose binding site. While inhibition and cell-based infection assays were beyond the scope of this work, the structural and solution-state data presented here establish a molecular framework for HMO recognition that will support future studies of their functional and antiviral properties and guide the design of antiviral compounds based on HMO architectures as classical methods such as ITC or BLI are not suitable for evaluation of ligand for HuNoVs.

## MATERIALS AND METHODS

### Ligands

Natural human milk oligosaccharides (HMOs) were provided by Glycom A/S, part of dsm-firmenich, and dissolved in H₂O to prepare stock solutions for experiments.

### Protein Expression and purification

The P domain (residues 224–530) of norovirus genogroup II variants GII.2/US/2022 (GenBank: OP689558.1), GII.4/Sydney/2016 (GenBank: MK756038.1) and GII.17/Rotterdam/2024 (GenBank: PQ336944.1) was recombinantly expressed in *Escherichia coli* BL21*(DE3) as an N-terminal His₆–MBP fusion incorporating a TEV protease cleavage site, using the pBADM-41 expression vector. Cells were cultured in LB medium supplemented with 100 µg/mL of ampicillin at 37 °C to an optical density at 600 nm of approximately 1.0 prior induction with 0.2% (w/v) L-arabinose and growth overnight at 16 °C. Harvested cells were resuspended in lysis buffer (20 mM Tris-HCl pH 8.0, 300 mM NaCl and 20 mM imidazole) using 5 mL per gram of cells and disrupted by sonication using a medium probe (3 cycles of 30 s at 60% amplitude with 0.5 s pulse intervals, with 2 min rest on ice between cycles). The fusion protein was captured by immobilized metal affinity chromatography using HisTrap HP column (Cytiva). The His₆–MBP tag was removed by overnight incubation at 20°C with homemade TEV protease at a 1/100 ratio prior to depletion of the MBP tag by amylose affinity chromatography (New England Biolabs). The P domain was subsequently polished by size-exclusion chromatography (SEC) on a TSKgel G2000SW column (Tosoh) equilibrated in 20 mM HEPES pH 7.4 and 150 mM NaCl. The dimeric state of the purified protein in solution was verified by SEC coupled to multi-angle light scattering (SEC-MALS) using a DAWN 8 detector (Wyatt Technology). After SDS-PAGE analysis, fractions containing purified proteins were pooled and concentrated to 3.5–5.5 mg/mL using centrifugal devices with 10 kDa cut-off PES membranes (Sigma-Aldrich, Merck) and stored at 4°C or flash-frozen in liquid nitrogen for storage at −80°C.

### Saturation Transfer Difference NMR and Binding Epitope Mapping

STD NMR (standard pulse sequence stddiff.3) was performed with a Bruker Avance III 500 MHz equipped with a cryoprobe TCI and connected to a Bruker SampleCase Plus autosampler. Spectra were recorded with Topspin 3.7.0. The proteins of the three different genotypes were exchanged into D_2_O PBS pH 7.4, the ligands were three times lyophilized against D_2_O. Each sample contained 15 µM of one of the three P-dimer genotypes, with 1500 µM HMO ligand. Each sample was measured at 5 ℃ degrees, with two different saturation times (t_sat_) of 0.75 s and 6 s, off saturation was set at 40 ppm with the on saturation at −1 ppm and 256 scans. The two different t_sat_ allows the use of the reduced dataset STD NMR (rd-STD NMR) approach for the generation of ligand binding epitope maps. Peak integrals were measured manually in TopSpin 4.7.0,

The saturation pulse had a strength of 40 dB. Binding epitope maps were generated with reduced dataset method described here. NMR spectra were processed with TopSpin v4.5.0 (Bruker) and MestReNova (version 16.0.0-39276).

### Crystallization and X-ray crystallography

Co-crystallization experiments were performed with a panel of HMOs, namely LNT, LNnT, 2’FL, 3FL LNFP-I/II/III/V, LNDFH-I/III, DFL and FSL. All crystals were grown exclusively by co-crystallization where each HMO was added at a final concentration of 6.5 mM to the purified protein at 3.5 mg/mL at least leading to a minimal molar ratio protein:ligand of 1:65. Hanging-drop vapor diffusion experiments were set up on the mix at 20 °C. Crystals of the GII.2 P domain (3.5 mg/mL) were obtained in reservoir solutions containing 20–22% PEG Smear Medium (Molecular Dimensions Ltd) supplemented with either 0.1 M sodium acetate or 0.05 M ammonium nitrate. The GII.4 P domain (3.5–5.0 mg/mL) crystallized in 18–20% PEG 3350 with 0.2 M sodium thiocyanate. The GII.17 P domain (3.5–5.0 mg/mL) crystallized in 14–16% PEG 3350 or PEG Smear Medium (Molecular Dimensions), 0.2 M magnesium sulfate or magnesium acetate and 0.1 M Tris-HCl pH 8.5. Prior to data collection, single crystals were cryoprotected by brief transfer into reservoir solution supplemented with 30% PEG and 20 mM of the respective HMO, then mounted in a cryoloop and flash-frozen in liquid nitrogen.

X-ray diffraction data were collected at 100 K on the Proxima 1 beamline at Synchrotron SOLEIL (Saint-Aubin, France) at a wavelength of 0.97856 Å using an Eiger X 16M detector (Dectris). Diffraction images were indexed, integrated and scaled using XDS and XDSME (40, 41), and downstream processing was performed with programs from the CCP4 suite v9.0 (42). Structures were determined by molecular replacement using Phaser (43), with previously determined P domain dimers structures as search models: PDB entry 4RPB for GII.2, 4OOV for GII.4 and 5ZUS for GII.17. Iterative cycles of maximum-likelihood refinement were carried out in REFMAC5 (44), interspersed with manual model rebuilding in Coot (45). Five percent of reflections were withheld throughout refinement for *R*_free_ calculation. HMOs ligands were modeled into unambiguous 2Fo–Fc and Fo–Fc electron density maps using standard carbohydrate monomer library (residue codes: FUC, GAL, NAG, BGC, SIA). Carbohydrate geometry, including ring conformation, regiochemistry and stereochemistry, was validated using Privateer (46, 47). Final model quality was assessed using the wwPDB validation server. Data collection and refinement statistics are reported in **Table S1 and S2**. Molecular graphics and figures were prepared with PyMOL Molecular Graphic System program (Version 2.0.4, Schrodinger, LLC). Protein–ligand interactions were initially identified using PLIP (Protein–Ligand Interaction Profiler) (48) and subsequently verified manually in PyMOL, with all interaction distances measured directly and contacts outside accepted geometric thresholds excluded from the final analysis. Buried surface areas were calculated using PISA (49). Atomic coordinates and structure factors for all seven structures have been deposited in the Protein Data Bank (PDB) under accession codes 28NP, 28SQ, 28UT, 28WR, 28ZE, 29AG and 29BS (50). Data collection and refinement statistics are reported in **Tables S2** and **S3**.

### Isotopically enriched Norovirus GII.4 Sydney P domain expression and purification

Uniformly ¹⁵N-labeled and ^2^H/^13^C/^15^N Norovirus GII.4/Sydney/2016 P domains were expressed in *E. coli* BL21 Gold (DE3) cells using the plasmid described above. For the production of uniformly ¹⁵N-labelled protein, overnight LB cultures were subsequently used to inoculate 1 L of M9 minimal medium containing ¹⁵NH₄Cl (1 g/L) as the sole nitrogen source, D-glucose (4 g/L), 1 mM MgSO₄, 0.1 mM CaCl₂, 1× vitamin mix, biotin solution (1 g/L) and ampicillin 100 µg/mL. To produce ^2^H/^13^C/^15^N - labelled protein, cells were grown in commercially available triple-labelling enriched medium (Silantes), according to the manufacturer’s recommendations. Cells were grown at 37 °C with shaking (180 rpm) to an OD₆₀₀ of 1.0-1.2, after which protein expression was induced with 0.2% (w/v) L-arabinose, followed by overnight incubation at 16 °C. Cells were harvested by centrifugation (7,500 ×g, 20 min, 4 °C) and resuspended in lysis buffer (20 mM Tris-HCl, 200 mM NaCl and 20 mM imidazole, pH 8.0) supplemented with protease inhibitors. Cell lysis was performed by sonication and lysates were clarified by centrifugation (35,000 ×g, 40 min, 4 °C). The soluble fraction was purified and concentrated as described above for the proteins in natural abundance. The only difference was that the size-exclusion chromatography (SEC) was performed using a HiLoad 16/600 Superdex 200 column (Cytiva) equilibrated with 20 mM Tris-HCl, 150 mM NaCl, pH 7.2.

### NMR Backbone assignment

NMR spectra for protein backbone assignment were acquired on 1200, 950, 900 and 500 MHz spectrometers (Bruker) equipped with a cryogenic probe at 310 K using TROSY versions of standard triple-resonance experiments, including HNCA, HNCO, HNCACB, and HN(CO)CACB spectra. ^2^H/^13^C/¹⁵N-labelled norovirus GII.4/Sydney/2016 P domain samples for NMR studies were prepared at a final concentration of 300 µM in NMR buffer consisting of 20 mM Tris, 150 mM NaCl, pH 7.2, 0.02% NaN₃, 0.05 mM EDTA and 1× protease inhibitor cocktail, supplemented with 10% (v/v) D₂O. Spectra were processed using Topspin 4.5.0 and analyzed with CARA (Computer Aided Resonance Assignment) software (51).

### Chemical shift perturbation (CSP) experiments

Ligand-binding experiments were performed using ¹H-¹⁵N TROSY-HSQC spectra recorded under the same buffer and temperature conditions described above for the resonance assignment (52). Samples of ¹⁵N-enriched norovirus GII.4 Sydney P domains were prepared at a concentration of 100 µM. These samples were then titrated by the stepwise addition of stock solutions of 2’FL, 3-FL, FSL and LNFP-I to achieve final ligand concentrations of 1, 4, 16, 32, 64 and 128 mM. Changes in cross-peak chemical shifts were monitored throughout the titration and the assigned resonances were used to quantify residue-specific chemical shift perturbations (CSPs). CSPs were calculated using Equation 1^39^ where Δδ_H_ and Δδ_N_ correspond to the chemical shift changes in the ¹H and ¹⁵N dimensions (in ppm), respectively.

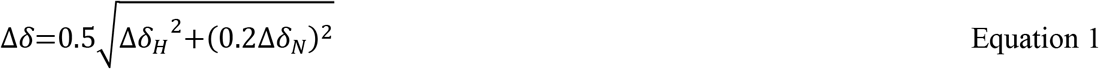

Residues with CSPs exceeding the mean plus one standard deviation (mean + σ) were classified as significantly perturbed. Dissociation constants (K_d_) were obtained by fitting CSP changes of selected residues to a one-site binding model (equation 2) where Δδ_obs_ is the observed CSP at each ligand concentration, Δδ_max_ is the maximum CSP at saturation, [P₀] is the total protein concentration, [L₀] is the total ligand concentration and K_d_ is the equilibrium dissociation constant (53). Nonlinear least-squares fitting was performed using OriginLab (OriginPro) software. The reported K_d_ represents the mean ± standard deviation (σ) derived from global fitting of well-resolved, significantly perturbed residues. For the ligands 2’FL and 3FL, apo, FSL and LNFP-I, the residues exhibiting significant chemical shift perturbations (CSPs) were color-coded according to the magnitude of the CSPs and mapped onto the surface representation of the crystal structure of the P domain to visualize the ligand-binding interface.

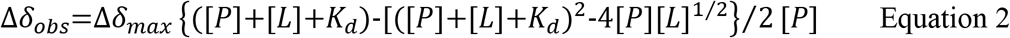

Δ*δ*: chemical shift; K_d_: binding affinity; [P]: protein concentration; [L]: ligand concentration

## ACKNOWLEDGMENTS

The HMOs were donated by Glycom A/S, part of dsm-firmenich, as a collaboration partner within the GlycoNoVi project. We thank Instruct-ERIC, and in particular the CERM/CIRMMP Italy center at the University of Florence, for access to the NMR facilities. The ICMG (UAR 2607) is acknowledged for providing access and support to the Octet RED96 Instrument on the PCI platform for BLI experiments. The authors would like to thank the synchrotron SOLEIL (Saint Aubin, France) for access to beamlines Proxima 1 (Proposal number 20231127) and technical support from the macromolecular crystallography staff. LP, AI and AV thanks CBH-EUR-GS (ANR-17-EURE-0003) and Glyco@Alps (ANR-15-IDEX-02) for support.

This work was supported by the European Union’s Horizon Europe research and innovation programme under the Marie Skłodowska-Curie grant agreement No. 101072717 (HORIZON-MSCA-2021-DN-01).

## AUTHOR CONTRIBUTIONS

Conceptualization: AI, AV, JA and MF; Formal analysis: AV, JCMG, LC, LP, HNF, RC, and SL; Funding acquisition: AI, JA and MF; Investigation: HNF, LC, LP, RC and SL; Project administration: AI, AV, JA and MF; Resources: LP and SL; Supervision: AI, AV, JA, JCMG and MF; Writing – original draft: LP, HNF and SL wrote the manuscript and Writing – review & editing: all authors have reviewed and agreed to the published version of the manuscript.

## DATA AVAILABILITY

The data that support the findings of this study are available in the Supporting Information of this article. The crystallographic data comprising coordinates and structure factors are publicly available from the PDB website using the ID codes provided in Table 1.

