## Supplemental tables and figures for "Structural Basis for Human Milk Oligosaccharides Recognition by Evolutionarily Dominant Norovirus Genogroup II Genotypes"

[e] Consorzio Interuniversitario di Risonanze Magnetiche di Metalloproteine (CIRMMP), 50019 Sesto Fiorentino, Italy.

§ Contributed equally

#### TABLE OF CONTENTS

##### List of Tables

**Table S1.** Epitope mapping of 2'FL, LNFP-III and LNDFH-III, elucidated with the reduced dataset STD-NMR approach ........................................................................S2

**Table S2.** Data collection and refinement statistics for GII.4 P domain complexes with human milk oligosaccharides ........................................................................S3

**Table S3.** Data collection and refinement statistics for GII.17 P domain complexes with human milk oligosaccharides ........................................................................S4

##### List of Figures

**Figure S1.** Proton assignment of the HMOs 2'FL, LNFP-III and LNDFH-III.......................S5

**Figure S2.** Electron density maps of HMO ligands bound to the GII.4 P domain..................S6

**Figure S3.** Electron density maps of HMO ligands bound to the GII.17 P domain................S6

**Figure S4.** Superposition of the GII.17 P domain in complex with α2Fuc bound ligands ....S7

**Figure S5. O**verlay of P domain overall structure among the three genotypes ......................S8

**Figure S6.** Global fitting CSP titration curves for HMO binding to the GII.4 P domain .......S9

**Figure S7.** Comparison of glycan-induced chemical shift perturbations with defined binding-site residues by crystallography in the GII.4 Sydney P domain ...............................S10

#### β-LNFP-III Epitope Map Table

| Proton Name | STD at short saturation time [s] | STD at long saturation time [s] | $k_{sat}$ | $STD_0$ | Epitope [%] | 20 uM GII.4 w/ 2 mM LNFP-III, 279K, tsat: 0.75 - 6 sec, -3ppm, 1024 scans, 700 MHz |
|---|---|---|---|---|---|---|
| Glc H1 α | 1.49 | 8.94 | | n.d. | | |
| Glc H1 β | 0.30 | 0.90 | 0.55 | 0.49 | 29.62 | |
| Glc H2 β | 0.379* | 0.60 | 1.32 | 0.79 | 47.91 | |
| Gal H1 | 0.33 | 0.85 | 0.66 | 0.56 | 33.60 | |
| Gal H4 | 0.26 | 0.76 | 0.55 | 0.42 | 25.10 | |
| GlcNAc H1 | 0.57 | 1.75 | 0.53 | 0.92 | 55.47 | |
| GlcNAc $CH_3$ | 0.72 | 3.55 | 0.30 | 1.07 | 64.60 | |
| Gal' H1 | 0.43 | 1.32 | 0.53 | 0.70 | 42.31 | |
| Gal ' H2 | 0.39 | 1.51 | 0.40 | 0.60 | 36.23 | |
| Fuc H1 | 0.80 | 5.21 | 0.22 | 1.17 | 70.23 | |
| Fuc H5 | 0.45 | 1.33 | 0.55 | 0.73 | 43.77 | |
| Fuc H6 | 0.85 | 1.52 | 1.09 | 1.66 | 100.00 | |

#### α-LNDFH-III Epitope Map Table

| Proton Name | STD at short saturation time [s] | STD at long saturation time [s] | $k_{sat}$ | $STD_0$ | Epitope [%] | 20 uM GII.4 w/ 2 mM LNDFH-III, 279K, tsat: 0.75 - 6 sec, -1/40ppm, 124 scans, 600 MHz |
|---|---|---|---|---|---|---|
| Glc H1 α | 2.18 | 8.02 | 0.42 | 3.39 | 90.30 | |
| Glc H1 β | n.d. | 1.95 | | | | |
| Glc H2 β | n.d. | n.d. | | n.d. | | |
| Fuc H1 α | 1.52 | 5.14 | 0.47 | 2.41 | 64.15 | |
| Fuc H1 β | 0.00 | 1.14 | | n.d | | |
| Fuc H5 | 0.00 | 2.5* | | | | |
| Fuc H6 | 1.68* | 2.5* | 1.50 | 3.76 | 100.00 | |
| Gal H1 | 0.42 | 2.35 | 0.26 | 0.62 | 16.52 | |
| Gal H3 | 0.94 | 2.71 | 0.57 | 1.53 | 40.72 | |
| GlcNAc H1 | 0.00 | 1.95 | | n.d. | | |
| GlcNAc $CH_3$ | 1.63 | 5.94 | 0.43 | 2.53 | 67.39 | |
| Gal' H1 | 0.37 | 2.19 | 0.25 | 0.54 | 14.31 | |
| Gal' H2 | * | * | | | | |
| Gal' H3 | * | * | | | | |
| Gal' H4 | 0.36 | 1.48 | 0.37 | 0.55 | 14.71 | |
| Fuc' H1 | 1.95 | 6.89 | 0.44 | 3.06 | 81.56 | |
| Fuc' H5 | n.d. | 2.5* | | n.d. | | |
| Fuc' H6 | 1.69 | 2.497* | 1.50 | 3.76 | 100.00 | |

#### β-2'FL Epitope Map Table

| Proton Name | STD at short saturation time [s] | STD at long saturation time [s] | $k_{sat}$ | $STD_0$ | Epitope [%] | 20 uM bs GII.4 with 2 mM 2'FL, 279K, tsat: 0.75 - 6 sec, 8.2ppm, 128 scans, 600 MHz |
|---|---|---|---|---|---|---|
| Glc H1 β | 0.66 | 1.05 | 1.33 | 1.40 | 78.41 | |
| Glc H2 β | n.d. | 1.35 | | | | |
| Glc H2 β | n.d. | 1.52 | | n.d. | | |
| Glc H3 β | n.d. | 1.35 | | | | |
| Gal H1 | 0.54 | 1.02 | 1.01 | 1.03 | 58.01 | |
| Fuc H1 | 1.19 | 5.59 | 0.32 | 1.78 | 100.00 | |
| Fuc H6 | 0.63 | 1.41 | 0.80 | 1.12 | 62.90 | |

**Table S1. Epitope Mapping of 2'FL, LNFP-III and LNDFH-III, elucidated with the reduced dataset approach.** STD intensities were measured by manual integration for short saturation time (0.75 s) and long saturation time (6 s). Details to the experiment are presented in the respective table (top right)

| Complex | GII.4 – FSL | GII.4 – LNFP-I | GII.4 – LNFP-III |
|---|---|---|---|
| ***Data collection*** | | | |
| Beamline | Soleil PX1 | Soleil PX1 | Soleil PX1 |
| Wavelength (Å) | 0.97856 | 0.97856 | 0.97856 |
| Space group | I 2 2 2 | I 2 | I 2 2 2 |
| Cell parameters – a, b, c (Å)<br>α, β, γ (°) | 86.92, 100.23, 130.48<br>90.00, 90.00, 90.00 | 86.58, 128.20, 102.70<br>90.00, 93.08, 90.00 | 87.29, 102.04, 130.34<br>90.00, 90.00, 90.00 |
| Protomers in asu | 2 | 4 | 2 |
| Resolution (Å)[a] | 46.28–1.95 (2.00–1.95) | 46.62–2.00 (2.04–2.00) | 46.49–2.60 (2.72–2.60) |
| No. reflections[a] | 363,438 (25,810) | 450,190 (26,886) | 155,232 (19,007) |
| No. unique reflections[a] | 41,854 (2,904) | 75,252 (4,432) | 17,655 (2,170) |
| CC1/2 (%)[a] | 99.8 (82.6) | 99.7 (79.5) | 99.3 (73.2) |
| Rmerge (within I+/I−)[a] | 0.104 (0.640) | 0.096 (0.621) | 0.162 (0.737) |
| Rmeas (within I+/I−)[a] | 0.118 (0.725) | 0.117 (0.753) | 0.181 (0.825) |
| Rpim (within I+/I−)[a] | 0.039 (0.240) | 0.066 (0.422) | 0.078 (0.361) |
| Mean I/σ(I)[a] | 12.8 (3.1) | 10.4 (2.4) | 9.6 (2.7) |
| Completeness (%)[a] | 100.0 (99.9) | 99.9 (99.7) | 97.2 (98.9) |
| Multiplicity[a] | 8.7 (8.9) | 6.0 (6.1) | 8.8 (8.8) |
| Wilson B-factor (Å$^2$) | 18.3 | 28.4 | 36.8 |
| ***Refinement*** | | | |
| Resolution (Å) | 46.28–1.95 | 46.62–2.00 | 41.76–2.60 |
| No. reflections / free reflections | 39,773 / 2,079 | 71,392 / 3,857 | 16,747 / 907 |
| Rwork / Rfree (%) | 14.4 / 19.9 | 17.9 / 23.3 | 19.0 / 26.5 |
| R.m.s. bond lengths (Å) | 0.018 | 0.006 | 0.005 |
| R.m.s. bond angles (°) | 2.105 | 1.431 | 1.308 |
| R.m.s. chiral (Å$^3$) | 0.118 | 0.071 | 0.061 |
| Clashscore | 1.56 | 1.85 | 2.73 |
| Fo, Fc correlation | 0.97 | 0.96 | 0.94 |
| No. atoms / B-factor (Å$^2$) | 5,561 / 25.857 | 10,205 / 32.149 | 4,942 / 39.258 |
| Protein | 4,864 / 24.875 | 9,544 / 32.036 | 4,784 / 39.025 |
| Ligand | 53 / 37.558 | 130 / 40.269 | 105 / 56.619 |
| Waters | 624 / 31.953 | 521 / 32.098 | 53 / 25.921 |
| Ramachandran | | | |
| Allowed | 3% | 3% | 4% |
| Favored | 97% | 97% | 96% |
| Outliers | 0% | 0% | 0% |
| PDB code | 28NP | 28UT | 28SQ |

[a] Values in parentheses are for the highest-resolution shell.

**Table S2. Data collection and refinement statistics for GII.4 P domain complexes with human milk oligosaccharides.**

| Complex | GII.17 – LNDFH-I | GII.17 – DFL | GII.17 – LNFP-II | GII.17 – LNFP-I |
|---|---|---|---|---|
| ***Data collection*** | | | | |
| Beamline | Soleil PX1 | Soleil PX1 | Soleil PX1 | Soleil PX1 |
| Wavelength (Å) | 0.97856 | 0.97856 | 0.97856 | 0.97856 |
| Space group | $P\ 2_1$ | $P\ 2_1$ | $P\ 2_1$ | $P\ 2_1$ |
| Cell parameters – a, b, c (Å) α, β, γ (°) | 58.81, 85.92, 64.21 90.00, 105.91, 90.00 | 59.41, 86.16, 64.06 90.00, 106.19, 90.00 | 59.50, 86.27, 63.97 90.00, 106.08, 90.00 | 59.22, 85.51, 65.62 90.00, 106.23, 90.00 |
| Protomers in asu | 2 | 2 | 2 | 2 |
| Resolution (Å)[a] | 47.24–1.10 (1.12–1.10) | 42.74–1.43 (1.46–1.43) | 47.66–1.53 (1.56–1.53) | 47.35–1.71 (1.74–1.71) |
| No. reflections[a] | 1,556,045 (66,820) | 764,704 (33,415) | 580,307 (27,161) | 225,417 (5,650) |
| No. unique reflections[a] | 247,453 (12,232) | 113,608 (5,395) | 92,749 (4,494) | 64,492 (1,689) |
| CC1/2 (%)[a] | 99.9 (76.5) | 99.9 (94.7) | 99.9 (89.3) | 99.0 (64.3) |
| Rmerge (within I+/I−)[a] | 0.055 (0.831) | 0.046 (0.323) | 0.052 (0.444) | 0.093 (0.524) |
| Rmeas (within I+/I−)[a] | 0.065 (1.030) | 0.055 (0.389) | 0.063 (0.537) | 0.127 (0.689) |
| Rpim (within I+/I−)[a] | 0.035 (0.599) | 0.029 (0.214) | 0.035 (0.300) | 0.086 (0.443) |
| Mean I/σ(I)[a] | 13.9 (1.8) | 20.1 (3.9) | 16.7 (3.3) | 6.9 (1.9) |
| Completeness (%)[a] | 99.8 (99.5) | 99.7 (94.6) | 99.8 (97.1) | 95.6 (47.4) |
| Multiplicity[a] | 6.3 (5.5) | 6.7 (6.2) | 6.3 (6.0) | 3.5 (3.3) |
| Wilson B-factor ($Å^2$) | 8.4 | 10.7 | 16.4 | 16.3 |
| ***Refinement*** | | | | |
| Resolution (Å) | 42.96–1.10 | 42.78–1.43 | 42.78–1.53 | 47.40–1.71 |
| No. reflections / free reflections | 234,893 / 12,522 | 107,812 / 5,767 | 88,033 / 4,687 | 61,282 / 3,183 |
| Rwork / Rfree (%) | 13.8 / 16.1 | 12.9 / 15.9 | 15.9 / 18.8 | 16.6 / 20.2 |
| R.m.s. bond lengths (Å) | 0.012 | 0.009 | 0.011 | 0.008 |
| R.m.s. bond angles (°) | 1.698 | 1.575 | 1.761 | 1.551 |
| R.m.s. chiral ($Å^3$) | 0.101 | 0.088 | 0.097 | 0.083 |
| Clashscore | 1.12 | 1.66 | 1.05 | 1.25 |
| Fo, Fc correlation | 0.98 | 0.97 | 0.97 | 0.96 |
| No. atoms / B-factor ($Å^2$) | 6057/16.571 | 5928/18.465 | 5704/19.414 | 5821/21.672 |
| Protein | 4871/13.824 | 4830/15.927 | 4804/18.077 | 4814/20.251 |
| Ligand | 136/ 19.183 | 86/19.534 | 94/26.025 | 94/28.818 |
| Waters | 1050/28.974 | 1012/30.487 | 806/26.613 | 913/28.427 |
| Ramachandran | | | | |
| Allowed | 2% | 2% | 2% | 2% |
| Favored | 98% | 98% | 98% | 98% |
| Outliers | 0% | 0% | 0% | 0% |
| PDB code | 28WR | 28ZE | 29BS | 29AG |

[a] Values in parentheses are for the highest-resolution shell.

**Table S3. Data collection and refinement statistics for GII.17 P domain complexes with human milk oligosaccharides**

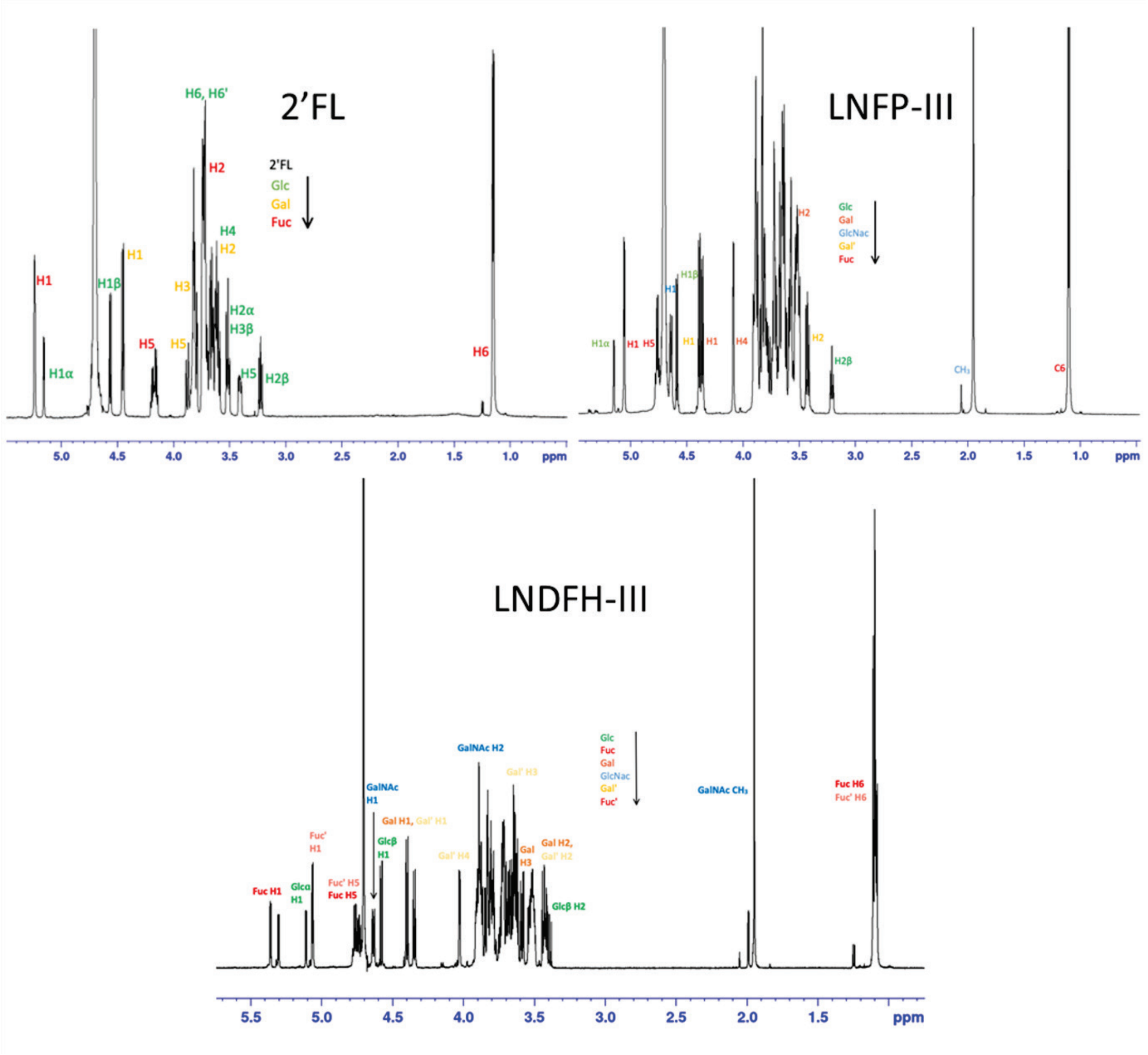


**Figure S1. Proton Assignment of the HMOs 2'FL, LNFP-III and LNDFH-III.** The protons were assigned with the help of Proton, COSY, DIPSY, NOESY and HSQC spectra with 2 mM of the HMO in $D_2O$.

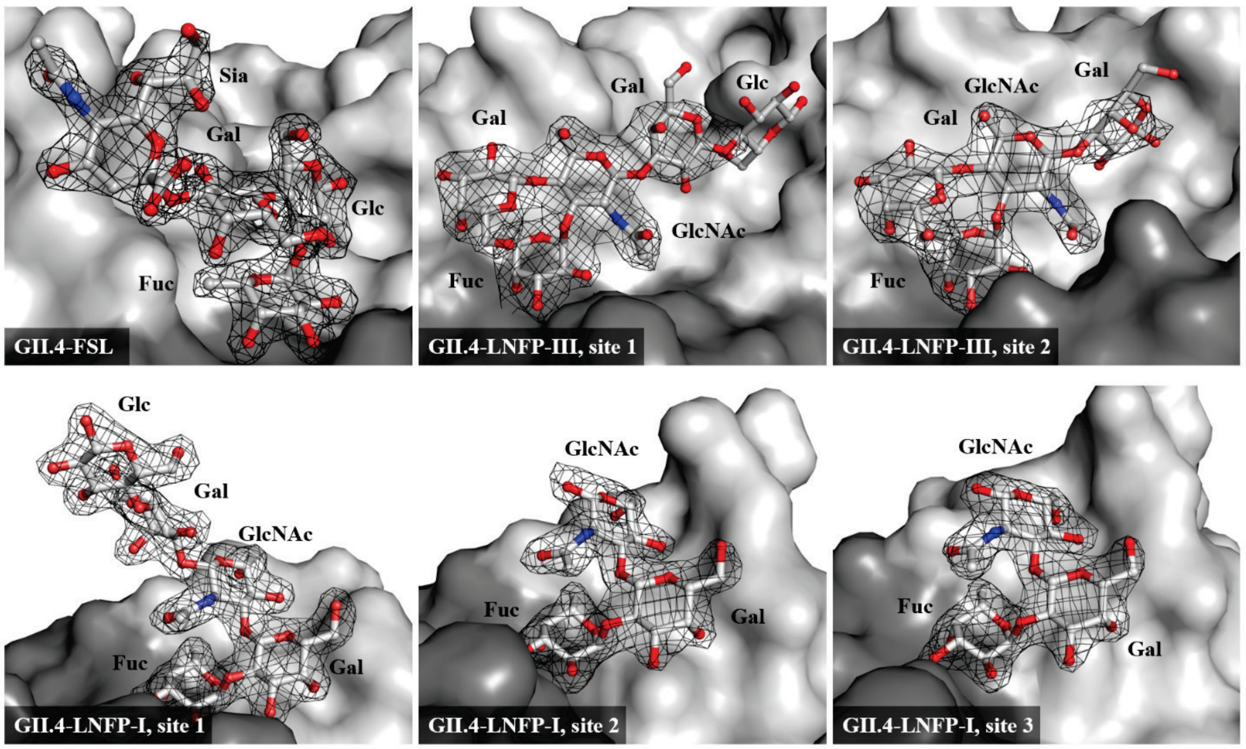


**Figure S2. Electron density maps of HMO ligands bound to the GII.4 P domain.** 2mFo-DFc electron density maps contoured at 1 sigma are shown around the bound sugars, on a surface representation of the protein in different shades of gray according to the protomer of the P dimer. For each ligand, all the binding sites observed in the asymmetric unit are shown: FSL, LNFP-III (sites 1 and 2) and LNFP-I (sites 1 to 3).

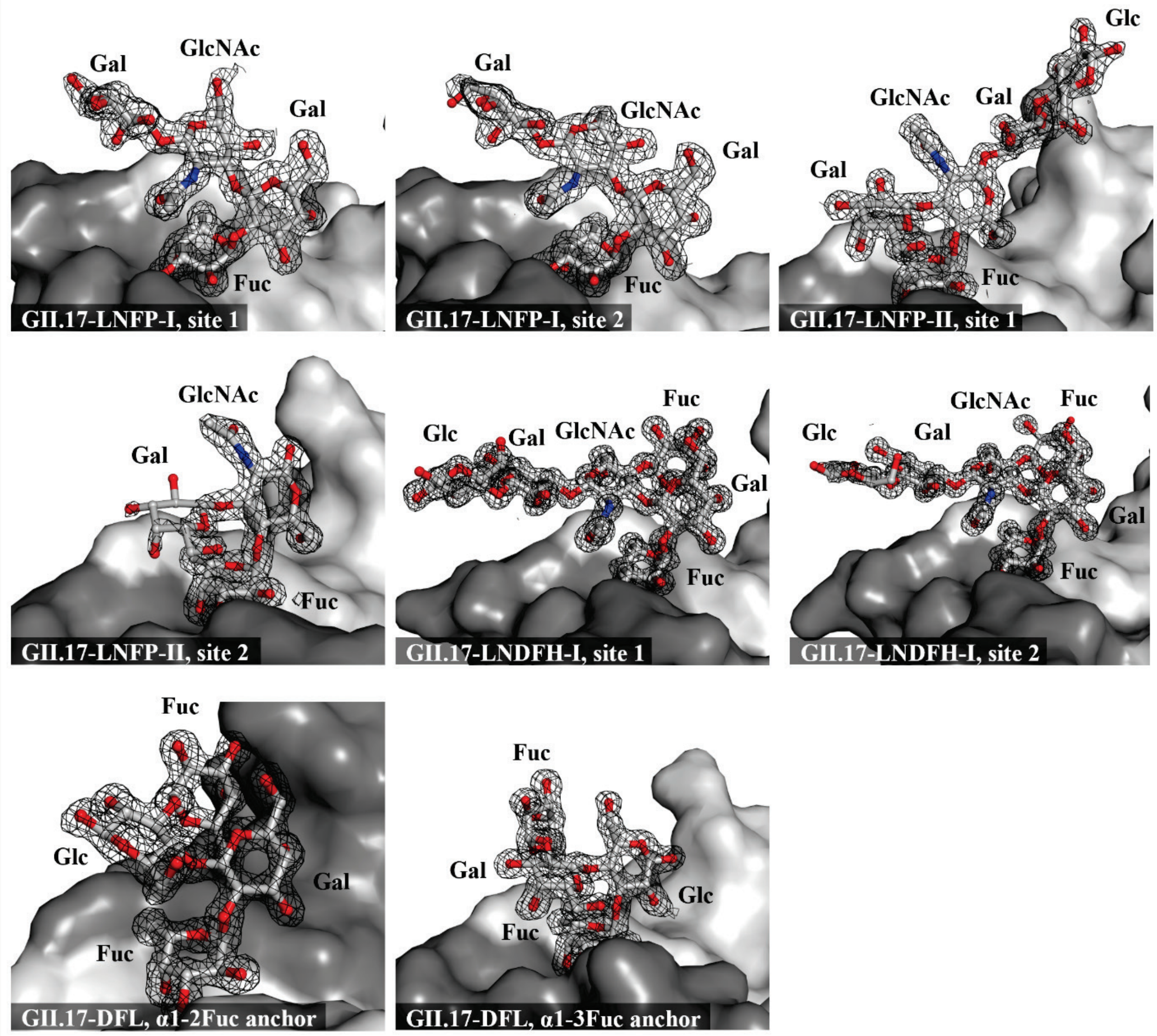


**Figure S3. Electron density maps of HMO ligands bound to the GII.17 P domain.** 2mFo-DFc electron density maps contoured at 1 sigma are shown around the bound sugars, on a surface representation of the protein in different shades of grey according to the protomer of the P dimer. For each ligand, all the binding sites observed in the asymmetric unit are shown: LNFP-I (sites 1 and 2), LNFP-II (sites 1 and 2), LNDFH-I (sites 1 and 2) and DFL (anchored through the alpha1-2 or the alpha1-3 linked fucose).

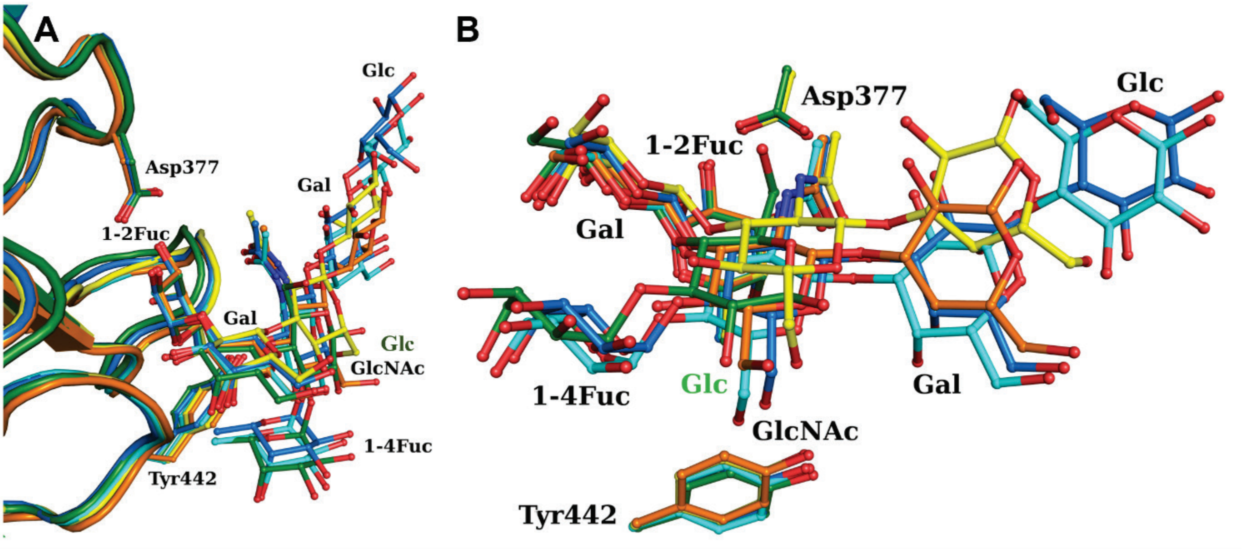


**Figure S4. Superposition of the GII.17 P domain in complex with α2Fuc bound ligands.** (**A**) Overall cartoon representation with ligand represented in balls and sticks for LNFP-I (PDB 29AG, yellow and orange), LNDFH-I (PDB 28WR, cyan and blue) and DFL (PDB 28ZE, dark green). (**B**) Focus on Tyr442 side chain and ligand changes in response to additional α4Fuc.

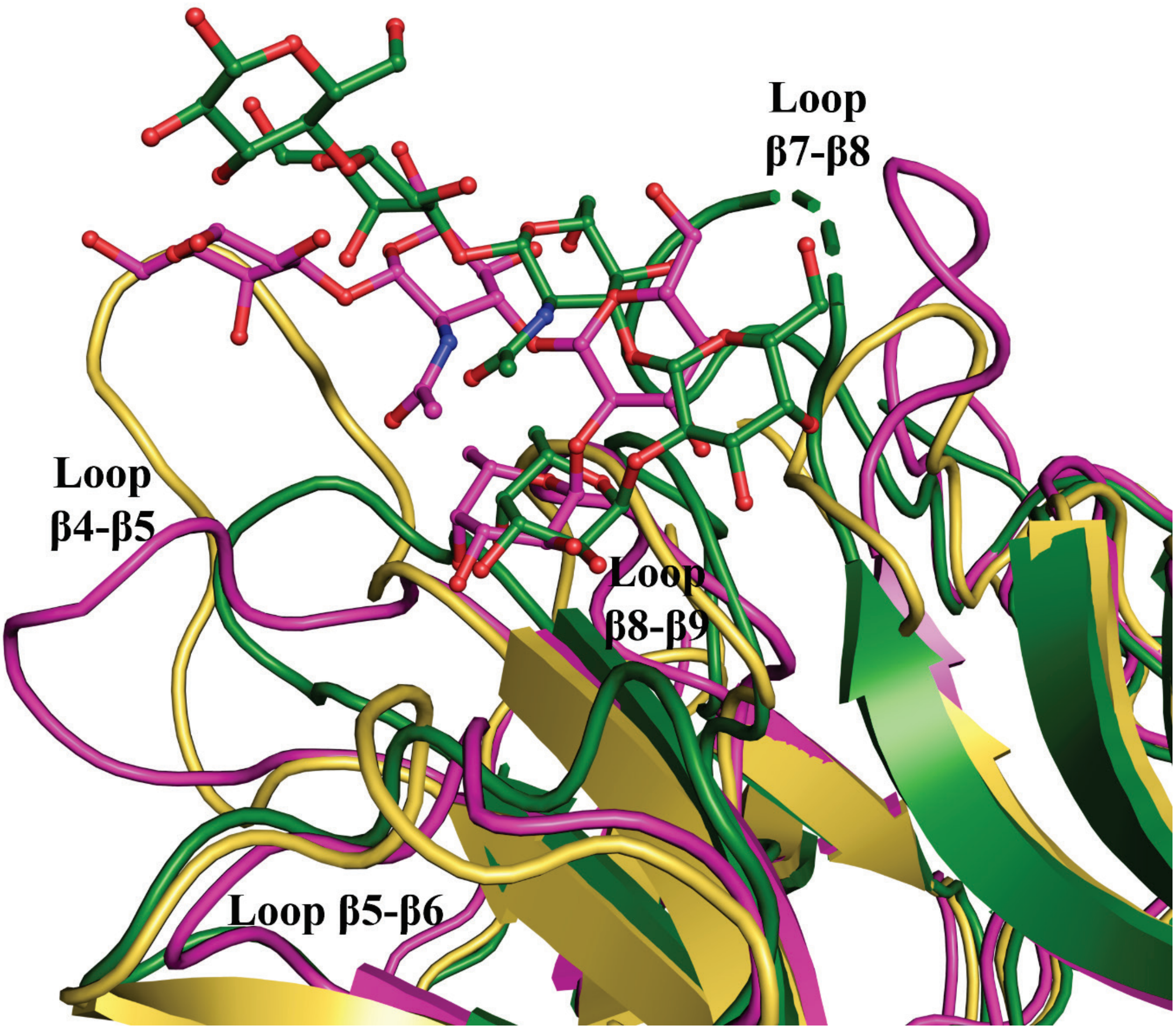


**Figure S5. Overlay of P domain overall structure among the three genotypes.** Superposition of the GII.4-LNFP-I (green), GII.17-LNFP-I (magenta) and GII.2 apo (yellow) with zoom on the surface loops around the binding site.

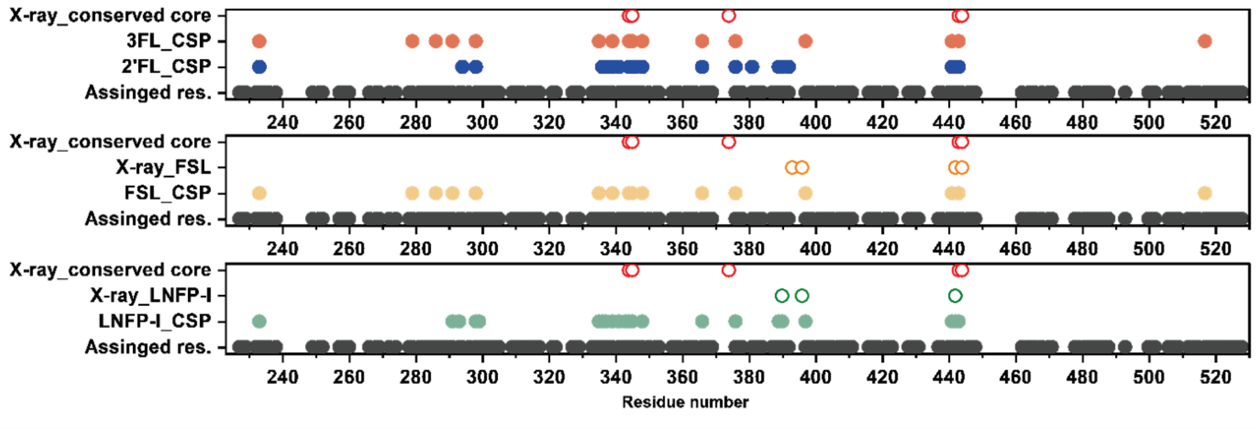


**Figure S6. Comparison of glycan-induced chemical shift perturbations with defined binding-site residues by crystallography in the GII.4 P domain.** Significant CSPs observed upon binding of 2′FL and 3FL, FSL and LNFP-I are plotted according to residue number. Filled coloured circles indicate residues showing significant CSPs for each ligand, whereas grey circles indicate backbone amide resonances assigned and therefore available for CSP analysis. Open red circles indicate residues belonging to the conserved crystallographic fucose-recognition site. Ligand-specific secondary contacts observed by X-ray crystallography are shown as open orange circles for FSL and open green circles for LNFP-I.

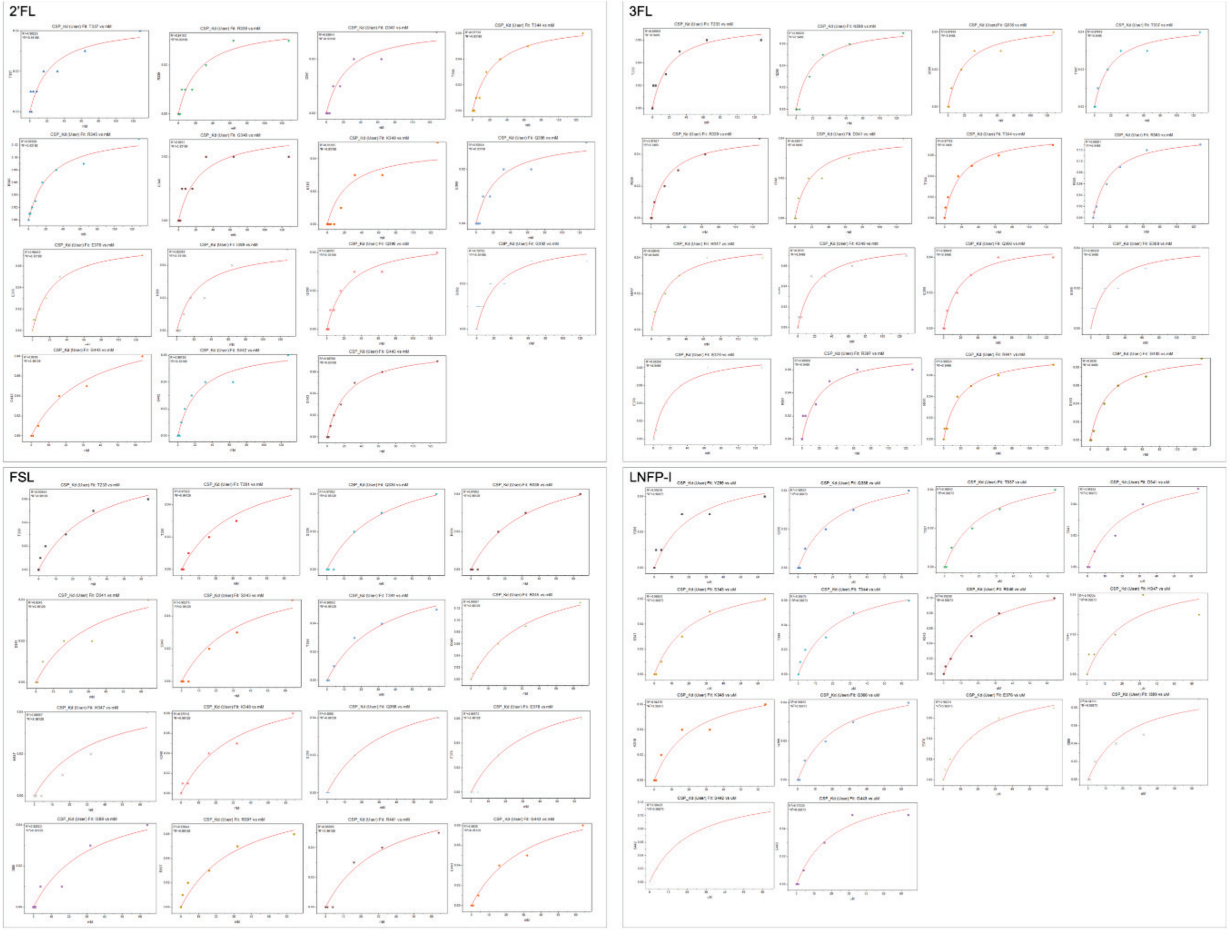


**Figure S7. Global fitting CSP titration curves for HMO binding to the GII.4 Sydney P domain.** Chemical shift perturbations for selected residues are plotted as a function of ligand concentration for 2′FL, 3FL, FSL and LNFP-I.
